# Deficiency in Galectin-3, -8, and -9 impairs immunity to chronic *Mycobacterium tuberculosis* infection but not acute infection with multiple intracellular pathogens

**DOI:** 10.1101/2022.12.29.522180

**Authors:** Huntly M. Morrison, Julia Craft, Rafael Rivera-Lugo, Jeffery R. Johnson, Guillaume R. Golovkine, Claire E. Dodd, Erik Van Dis, Wandy L. Beatty, Shally R. Margolis, Teresa Repasy, Isaac Shaker, Angus Y. Lee, Russell E. Vance, Sarah A. Stanley, Nevan J. Krogan, Dan A. Portnoy, Bennett H. Penn, Jeffery S. Cox

## Abstract

Macrophages employ an array of pattern recognition receptors to detect and eliminate intracellular pathogens that access the cytosol. The cytosolic carbohydrate sensors Galectin-3, -8, and -9 (Gal-3, Gal-8, and Gal-9) recognize damaged pathogen-containing phagosomes, and Gal-3 and Gal-8 are reported to restrict bacterial growth via autophagy in cultured cells. However, the contribution of these galectins to host resistance during bacterial infection remains unclear. We found that Gal-9 binds directly to *Mycobacterium tuberculosis* (*Mtb*) and *Salmonella enterica* serovar Typhimurium (*Stm*) and localizes to *Mtb* in macrophages. To determine the combined contribution of membrane damage-sensing galectins to immunity in vivo, we generated Gal-3, -8, and - 9 triple knockout (TKO) mice. *Mtb* infection of primary macrophages from TKO mice resulted in defective lysosomal trafficking but normal bacterial replication. Surprisingly, these mice had no discernable defect in resistance to acute infection with *Mtb*, *Stm or Listeria monocytogenes*, and had only modest impairments in bacterial growth restriction and CD4 T cell activation during chronic *Mtb* infection. Collectively, these findings indicate that while Gal-3, -8, and -9 respond to an array of intracellular pathogens, together these membrane damage-sensing galectins play a limited role in host resistance to bacterial infection.

**Author Summary:** Intracellular bacterial pathogens cause many of the world’s most deadly infectious diseases. A common requirement for nearly all intracellular pathogens is the ability to damage the endomembrane compartments in which they reside, which allows pathogens access to the nutrient-rich cytosol of the host. However, membrane damage also creates a “pattern of pathogenesis” that triggers antimicrobial immune responses. Galectin-3, -8, and -9 (Gal-3, Gal-8, and Gal-9) act as a surveillance system for membrane damage and Gal-3 and Gal-8 inhibit bacterial growth by activating autophagy, a cellular pathway that can capture cytosolic bacteria and degrade them in lysosomes. Membrane damage-sensing galectins were hypothesized to promote bacterial killing during acute infection yet their role in the immune response of an infected animal remains unclear. Here, we show that mice deficient for Gal-3, -8, and -9 had no defects in resistance to acute infection with the pathogens *Listeria monocytogenes, Salmonella enterica* serovar Typhimurium, and *Mycobacterium tuberculosis* (*Mtb*), and were only modestly susceptible to chronic *Mtb* infection. Our data suggest that Gal-3, -8 and -9 are not critical for innate immune responses during acute infection and may play a more prominent role in the adaptive immune response. These results broaden our understanding of the role of membrane damage-sensing pathways in host defense against bacterial infection.

## Introduction

Infectious diseases caused by intracellular bacterial pathogens are among the leading causes of mortality worldwide [1]. Bacterial replication within host cells is critical for pathogenesis yet also exposes bacteria to a myriad of innate immune sensing mechanisms [2]. Detection of pathogen-associated molecular patterns (PAMPs) by pattern recognition receptors (PRRs) activates multiple cell-intrinsic antimicrobial pathways, including autophagy and inflammasome assembly [3, 4], and proinflammatory signaling pathways required for coordinating protective adaptive immune responses [5, 6]. However, the relative importance and precise function of distinct pathogen-sensing pathways during in vivo infection remains poorly understood.

Macrophages can sense and respond to intracellular pathogens using galectins [7–9], a family of soluble β-galactoside-binding receptors that facilitate carbohydrate sensing in diverse biological contexts, including chronic inflammation, autoimmunity, cancer, and infection [10–12]. There are fifteen galectins in mammals, each containing one or two carbohydrate-recognition domains (CRDs) [12]. While some galectins are reported to act extracellularly through an atypical secretion mechanism [10, 13], a subset, Galectin-3, -8, and -9 (Gal-3, Gal-8, and Gal-9), act as sentinels for intracellular membrane damage [7, 9]. Lysosomal membrane damage activates Gal-3, -8, and -9, which promote repair and clearance of damaged lysosomes [14–18]. Several bacterial pathogens, including *Mycobacterium tuberculosis* (*Mtb*)*, Salmonella enterica* serovar Typhimurium (*Stm)*, *Listeria monocytogenes* (*Lm*), *Shigella flexneri*, and *Legionella pneumophila (Lp*), inflict phagosomal membrane damage, exposing luminal glycoproteins that recruit Gal-3, -8, and -9 via their CRDs [7–9,19–21]. Additionally, some galectins can interact directly with bacteria and their cell wall glycans [22–25]. In particular, Gal-9 was recently shown to bind *Mtb* via recognition of the cell wall polysaccharide, arabinogalactan [25]. During infection, galectin recruitment and signaling promote antibacterial autophagy and bacterial growth restriction [9,15,20], proinflammatory signaling [25], and recruitment of antibacterial guanylate-binding proteins [21]. In addition, we previously found that Gal-3, -8, and -9 are all ubiquitylated during *Mtb* infection [26, 27], although the implications of these modifications remain unclear.

Galectins activate autophagy, a conserved process that targets, captures, and degrades cytosolic cargo, including intracellular pathogens [28, 29]. Host cells employ several mechanisms to target cytosolic bacteria to the autophagy pathway, deploying ubiquitin ligases and PRRs, such as cGAS [3], and Gal-3 and Gal-8, which promote recruitment of autophagy proteins via distinct mechanisms [9,15,20]. *Mtb*-mediated perforation of the phagosome via the bacterial ESX-1 secretion system promotes recruitment of ubiquitin ligases Parkin and Smurf1, which initiate autophagy targeting by ubiquitylating substrates around bacteria with K63- and K48-linked polyubiquitin chains [30, 31].

Several ubiquitin-binding autophagy receptors, including SQSTM1, NDP52, TAX1BP1, NBR1, and OPTN, decode these ubiquitin signals into autophagy signaling cascades that nucleate autophagosome membranes around bacteria for trafficking to lysosomes [20,26,32,33]. Autophagy receptors are also recruited directly to pathogen-containing phagosomes via interactions with Gal-8, which has been shown to promote autophagic clearance of *Stm* and *Mtb* via recruitment of NDP52 and TAX1BP1, respectively [9, 20].

While Gal-3, -8, and -9 have each been shown to promote autophagy in cultured cells [9,15,18,20], the contribution of these membrane damage-sensing galectins to the host immune response in vivo remains incompletely understood. Following high-dose aerosol infection, mice lacking Gal-3 or Gal-8 succumbed slightly faster to *Mtb* infection [15, 16], but it remains unclear whether bacterial control or adaptive immune responses differ in mice lacking these galectins. Similarly, high-dose intranasal infection of Gal-9 knockout mice resulted in a modest 2-fold increase in *Mtb* burden [25]. The potential for redundancy between these galectins, which all respond to membrane damage, could explain the seemingly mild phenotypes of individual galectin knockouts. In addition, the roles of these galectins in murine models of *Lm* and *Stm* infection remain unexplored. Thus, overall, the roles of membrane damage-sensing galectins in host resistance to bacterial infection remain poorly understood.

In the present study, we identified Gal-9 in a mass spectrometry-based search for host proteins that bind to the surface of *Mtb*. We found that Gal-9 bound *Mtb* in a carbohydrate-dependent manner and, similar to Gal-3 and Gal-8, localized to bacteria in response to phagosome damage during macrophage infection. To further characterize the collective role of membrane damage-sensing galectins in vivo, we generated Gal-3, -8, and -9 triple knockout (TKO) mice. Primary bone marrow-derived macrophages (BMMs) from TKO mice exhibited mild impairment in *Mtb*-targeted autophagy yet maintained normal bacterial growth restriction during infection with multiple intracellular pathogens. Furthermore, TKO mice exhibited no defects in resistance to acute bacterial infection with *Mtb, Lm,* or *Stm*, and showed only modest susceptibility to chronic *Mtb* infection. These results suggest that membrane damage-sensing galectins play a limited role in antibacterial immunity in vivo and may fulfill roles other than mediating direct antibacterial resistance.

## Results

### Gal-9 binds to *Mtb*

To identify host proteins that bind to the *Mtb* surface, we performed pull-down experiments with formaldehyde-fixed *Mtb* incubated with lysates from differentiated THP-1 macrophages (Fig 1A). After washing the bacteria, the bound proteins were eluted with 8M urea and analyzed by liquid-chromatography mass spectrometry (LC-MS/MS) (Fig 1A). Of the ∼300 human proteins identified (Table S1), we focused on lipid- and carbohydrate-binding proteins, as these molecules are likely to recognize cell wall components during infection. Intriguingly, this dataset included the carbohydrate-binding lectins, Gal-1 and Gal-9. Since prior studies have shown that Gal-1 does not localize to bacterial phagosomes or appear to play a role in pathogen sensing [9, 20], we focused on Gal-9. We detected five unique peptides that mapped to Gal-9, which is composed of two distinct CRDs that are joined by a linker region (Fig 1B). To biochemically validate the LC-MS/MS results, we expressed FLAG-tagged mouse Gal-9 in RAW 264.7 mouse macrophages and performed similar pull-down experiments with formaldehyde-fixed *Mtb* as described above, followed by anti-FLAG immunoblotting. Consistent with our findings with human Gal-9 from THP-1 macrophages, mouse Gal-9 also bound to the surface of *Mtb* (Fig 1C). This interaction was dependent upon carbohydrate binding, as addition of lactose to the in vitro binding reaction blocked Gal-9 binding (Fig 1C). We used pull-down assays to further explore whether Gal-9 binds to other phylogenetically diverse intracellular pathogens and found that Gal-9 bound to *Stm* and did not bind to *Lm* or the fungal pathogen *Cryptococcus neoformans* (*Cn*) under the tested conditions (Fig 1C).

**Fig 1.**
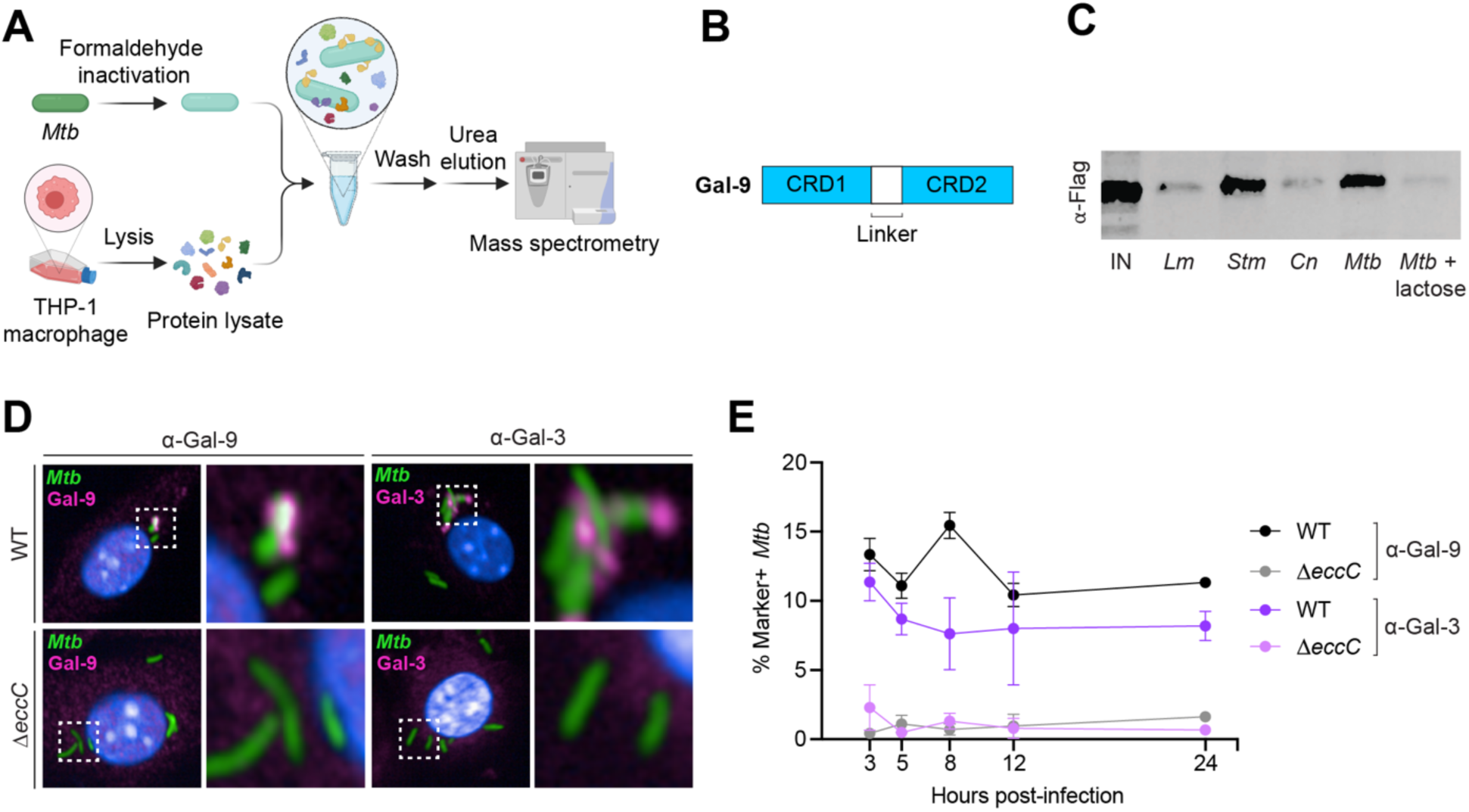
Gal-9 binds and recruits to *Mtb.* (**A**) Experimental design for *Mtb* pull-down mass spectrometry identification of *Mtb*-binding proteins. (**B**) Domain organization of Gal-9. CRD, carbohydrate recognition domain. (**C**) Immunoblot of in vitro binding reactions between indicated pathogens and Gal-9-FLAG THP-1 lysate, probed with anti-FLAG antibody; IN, input; *Lm*, *Listeria monocytogenes*; *Stm*, *Salmonella enteria* serovar Typhimurium; *Cn*, *Cryptococcus neoformans*; *Mtb*, *Mycobacterium tuberculosis*. (**D**) Confocal microscopy of WT BMMs infected with WT or Δ*eccC Mtb*-GFP (MOI = 2) 8 hours post-infection and immunostained for endogenous Gal-9 and Gal-3. (**E**) Quantification of *Mtb*-GFP colocalization with Gal-9 or Gal-3 at indicated time points. Figures represent two independent experiments (D, E). Error bars represent SD from 3 technical replicates.

To determine whether Gal-9 localizes to *Mtb* in infected macrophages, we analyzed *Mtb*-infected Gal-9-FLAG cells using immunogold electron microscopy with anti-FLAG antibody (Fig S1). Gal-9-FLAG was found primarily on extra-phagosomal membranes throughout the macrophage cytoplasm, with roughly 25% of puncta localized to phagosomal membranes and 5% localizing to bacteria (Fig S1). Taken together, these results demonstrate that Gal-9 can bind directly to *Mtb* via carbohydrate-binding and that Gal-9 localizes to *Mtb* in an infected macrophage.

### Gal-9 recruitment to *Mtb* is ESX-1-dependent

To determine whether ESX-1-mediated membrane damage is required for the recruitment of endogenous Gal-9, we infected BMMs with fluorescent *Mtb* and used automated confocal microscopy to quantify Gal-9 recruitment to >4000 bacteria. *Mtb*-Gal-9 colocalization was evident as early as 3 hours post-infection and reached a maximum of 15% at 8 hours post-infection before returning to 10-12% at later times (Fig 1D, 1E). We observed minimal colocalization between Gal-9 and the ESX-1 secretion mutant Δ*eccC*, indicating that Gal-9 recruitment requires ESX-1-mediated membrane damage and cytosolic access. As a comparison, we also measured endogenous Gal-3 recruitment to *Mtb* and found that Gal-3 similarly colocalized with approximately 8-11% of WT *Mtb* but showed no recruitment to Δ*eccC Mtb* (Fig 1D, 1E). Thus, endogenous Gal-9 is recruited to *Mtb* in an ESX-1-dependent manner soon after infection and maintains steady localization with a subpopulation of bacteria throughout infection.

### Generation and validation of Gal-3, -8, and -9 triple knockout (TKO) mice

While many cell-intrinsic roles have been described for Gal-3, -8, and -9, their contributions to host immunity are incompletely understood. Prior studies have shown that Gal-3, -8, and -9 each make partial contributions to *Mtb* immunity in animal models [15,16,25]. Because the mild phenotypes of individual galectin knockouts could be due to functional redundancy between damage-responsive galectins, we sought to test the combined role of all three membrane damage-sensing galectins in antibacterial immunity in vivo by generating Gal-3, -8, and -9 triple knockout (TKO) mice. Using CRISPR/Cas9 genome editing, we generated two independent TKO mouse strains (strain 1 and strain 2), both containing frameshift mutations in each of the three galectins that result in premature stop codons. Western blotting of BMMs from both TKO strains confirmed that these mutations led to loss of protein expression (Fig 2A, S2A). Gal-3, -8, and -9 have documented roles in the response to lysosomal damage, which entails activation of lysophagy and ESCRT-mediated repair [15–18]. To determine whether TKO cells have defective lysosomal repair responses, we treated WT and TKO BMMs with the lysosomal damaging agent Leu-Leu-OMe (LLOMe) and stained with Lysotracker to assess lysosome abundance and integrity. There was a modest reduction in lysotracker puncta per cell in TKO BMMs using two different treatment conditions (Fig S2B, S2C), indicating an impairment in lysosomal repair in galectin-deficient cells. Thus, we have established a genetic system to evaluate the collective roles of membrane damage-sensing galectins in vivo.

**Fig 2.**
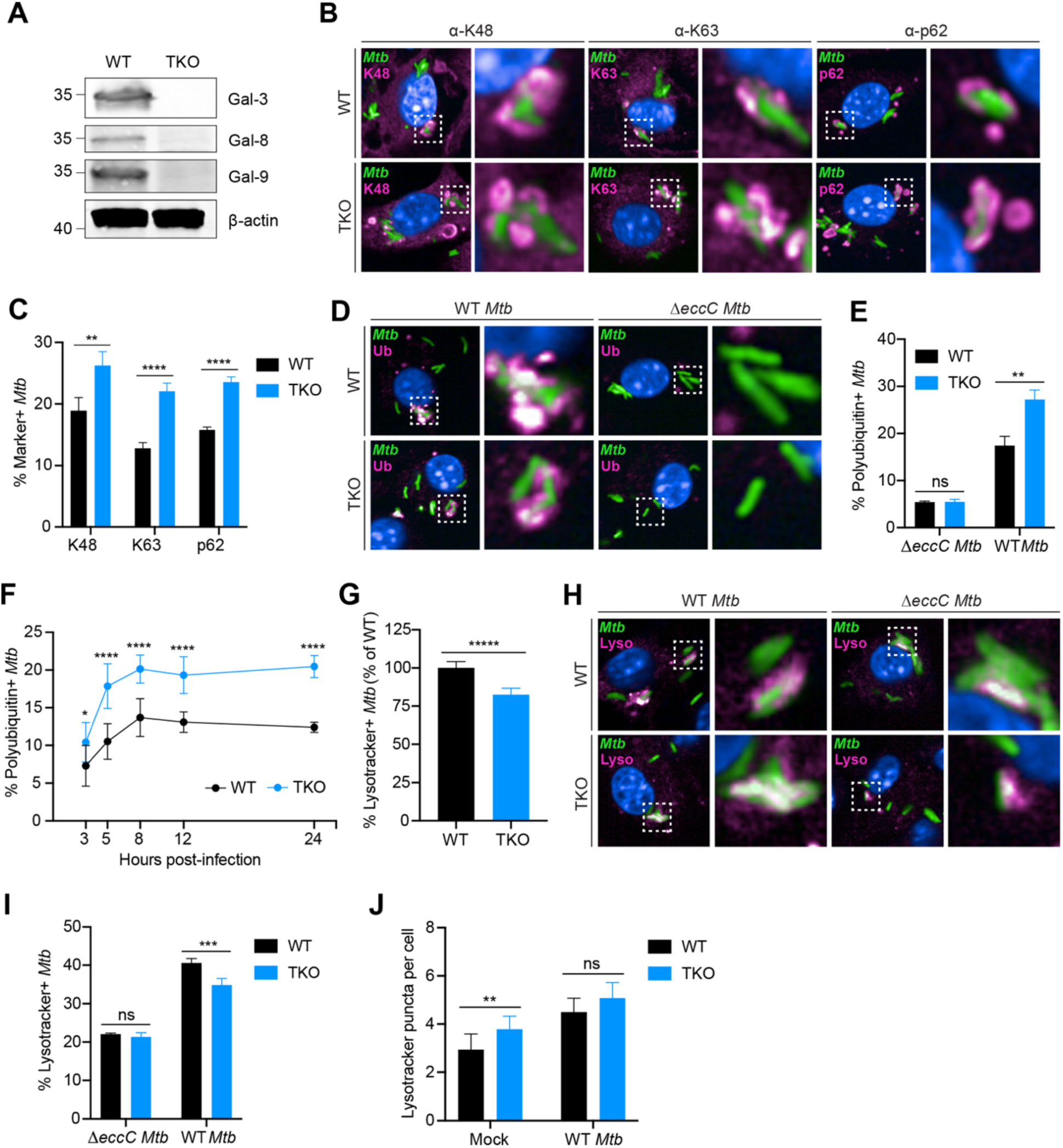
TKO macrophages exhibit impaired *Mtb* lysosomal trafficking and accumulate autophagy targeting intermediates. (**A**) Immunoblots of bone marrow-derived macrophage lysates from WT and TKO mice (strain 1) probed for indicated proteins. (**B**) Confocal microscopy of WT or TKO BMMs infected with *Mtb-*GFP (MOI = 2) 8 hours post-infection and immunostained for K48- or K63-linked polyubiquitin, or p62 as indicated. (**C**) Quantification of (B) for *Mtb-*GFP colocalization with indicated markers. (**D**) Same as in (B) but infected with WT or Δ*eccC Mtb*-GFP and immunostained for polyubiquitin. (**E**) Quantification of (D) for WT or Δ*eccC Mtb-*GFP colocalization with polyubiquitin. (**F**) Quantification of *Mtb-*GFP colocalization with polyubiquitin in WT or TKO BMMs at indicated time points. (**G**) Quantification of *Mtb*-GFP colocalization with Lysotracker signal. (**H**) Same as in (D) but stained with Lysotracker. (**I**) Quantification of (H) for WT or Δ*eccC Mtb* colocalization with Lysotracker signal. (**J**) Quantification of Lysotracker puncta 8 hours post-infection in mock- or *Mtb*-GFP-infected (MOI = 2) WT or TKO BMMs. Figures represent two (F, J), or three (G) independent experiments, or are representative of two independent experiments (B-E, H-I). Error bars represent SD from four technical replicates, and *p<0.05., **p<0.01, ***p<0.001, ****p<0.0001 by unpaired t-test.

### Deficiency in Gal-3, -8, and -9 leads to inefficient *Mtb* lysosomal trafficking and accumulation of autophagy targeting intermediates

Given the characterized roles of Gal-3 and Gal-8 in antibacterial autophagy induction [9,15,20], we tested whether TKO macrophages were impaired for *Mtb*-targeted autophagy, which is characterized by recruitment of polyubiquitin chains and ubiquitin-binding autophagy receptors to cytosolic *Mtb* [30, 33]. We used automated confocal microscopy to quantify the recruitment of polyubiquitin, polyubiquitin linkage subtypes K48 and K63, and p62 to *Mtb* in both WT and TKO BMMs. We hypothesized that decreased *Mtb*-ubiquitin colocalization would indicate an impairment of autophagy initiation, whereas an increase would indicate either a block in *Mtb* delivery to the lysosome or a defect in phagosome repair, which could expose bacteria to increased autophagy targeting. TKO macrophages exhibited a 40-70% increase in *Mtb* colocalization with all autophagy markers (Fig 2B-F). The increase in *Mtb*-ubiquitin colocalization persisted from 3 to 24 hours post-infection (Fig 2F), indicating a sustained block in either phagosomal repair or lysosomal trafficking of cytosolic *Mtb*. As expected, the increase in ubiquitin-labeled bacteria was dependent upon ESX-1-mediated phagosome permeabilization (Fig 2D, 2E). We also measured *Mtb* colocalization with TAX1BP1 and OPTN, which are autophagy receptors implicated in *Mtb*-targeted autophagy [20,26,33]. In contrast to p62, there were no differences in TAX1BP1 and OPTN recruitment to *Mtb* between WT and TKO macrophages (Fig S3).

To assess if autophagosome maturation was impaired, we measured *Mtb* delivery to the lysosome by infecting cells with *Mtb* and staining with Lysotracker. TKO macrophages showed a 20% decrease in *Mtb-*Lysotracker colocalization relative to WT (Fig 2G). This loss in colocalization was not due to reduced lysotracker staining from defects in lysosomal homeostasis or repair in TKO cells (as seen during LLOMe treatment in Fig S2B, S2C), as both mock- or *Mtb*-infected WT or TKO macrophages had similar numbers of Lysotracker puncta (Fig 2J). In contrast to WT *Mtb*, infection with *ΔeccC Mtb* resulted in similar levels of *Mtb-*Lysotracker colocalization in WT and TKO cells (Fig 2H, 2I), indicating that galectin-dependent lysosomal trafficking requires ESX-1 and cytosolic access. Taken together, these data suggest that while Gal-3, -8, and -9 are dispensable for the initiation of ubiquitin-mediated selective autophagy and p62 recruitment, they are necessary for efficient delivery of cytosolic *Mtb* to the lysosome.

### Gal-3, -8, and -9 are not required for control of intracellular bacterial pathogens in primary macrophages

We next tested whether TKO BMMs were more permissive for *Mtb* replication by measuring bacterial growth of a bioluminescent *Mtb* reporter strain (*Mtb*-LUX) [34]. We found that despite defects in lysosomal trafficking in TKO macrophages, bacterial growth kinetics were nearly identical between WT and TKO macrophages (Fig 3A, 3B), a finding we confirmed in separate experiments by directly enumerating *Mtb* colony-forming units (CFUs) (Fig 3C). In addition, TKO macrophages exhibited normal interferon-gamma (IFN-γ) dependent restriction of *Mtb* (Fig 3A). We also observed no differences in *Mtb* growth in two different Gal-9 shRNA knockdown THP-1 cell lines compared to control shRNA (Fig 3D), demonstrating that Gal-9 is nonessential for restriction of *Mtb* growth in immortalized human macrophages. Thus, despite previously described roles for Gal-3 and Gal-8 in autophagy and restriction of bacterial growth in immortalized cell lines [9,15,20], we found no defects in cell-intrinsic *Mtb* restriction in the absence of membrane damage-sensing galectins.

**Fig 3.**
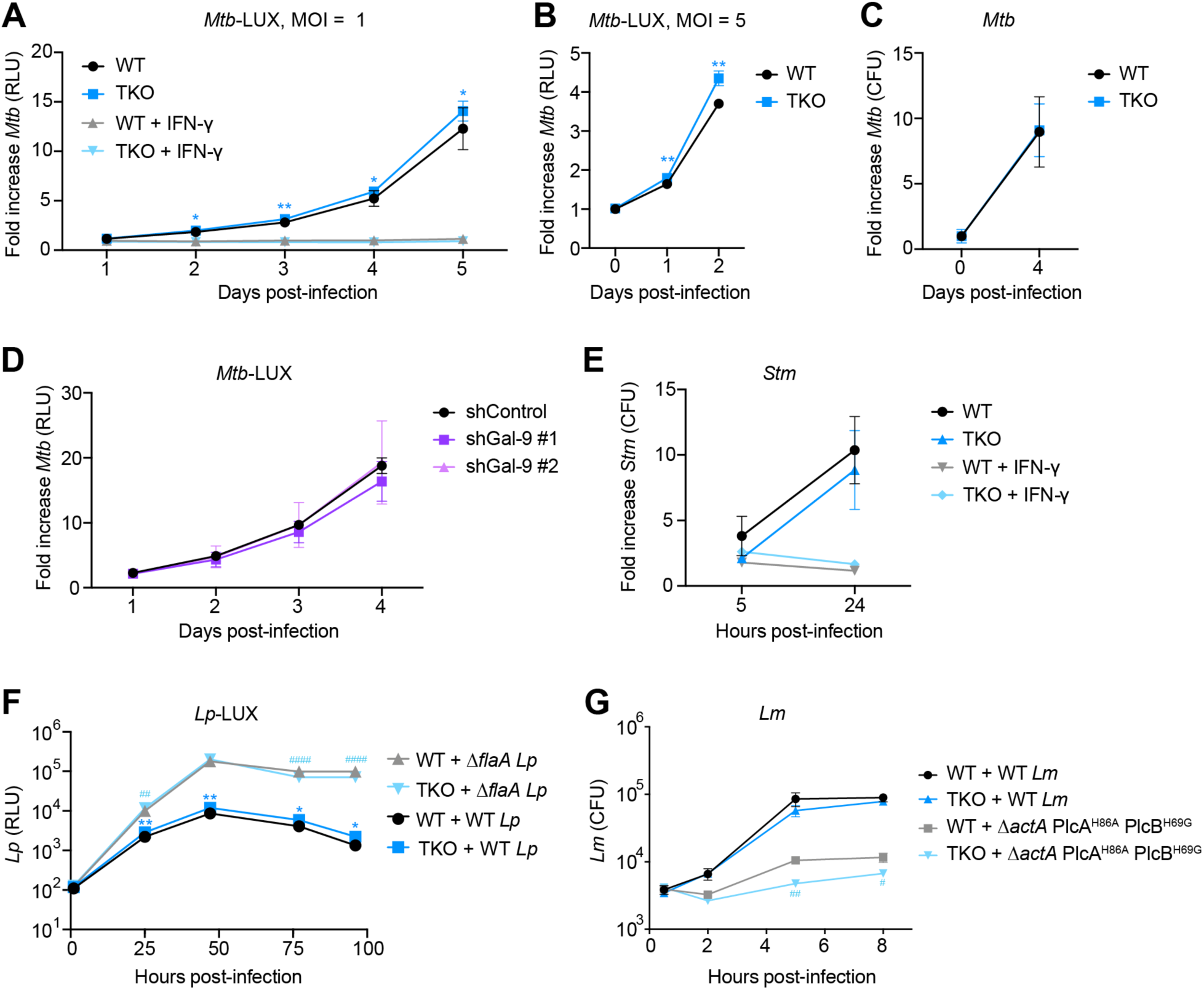
Gal-3, -8, and -9 are not required for bacterial growth restriction in macrophages. (**A and B**) Relative light units (RLU) fold change of WT or TKO BMMs infected with *Mtb-*LUX (MOI = 1, A; MOI = 5, B). (**C**) Colony-forming unit (CFU) fold change of WT or TKO BMMs infected with WT *Mtb* (MOI = 2). (**D**) RLU-fold change of control or Gal-9 knockdown shRNA THP-1 cells infected with *Mtb-*LUX (MOI = 1). (**E**) CFU-fold change of WT or TKO BMMs infected with WT *Stm* (MOI = 10). (**F**) RLU of WT or TKO BMMs infected with WT or Δ*flaA Lp-*LUX (MOI = 0.01). (**G**) CFU per coverslip of WT or TKO BMMs infected with WT or Δ*actA* PlcA^H86A^ PlcB^H69G^ *Lm* (MOI = 0.25). Figures represent one (B, F), two (G, A), or four (D) independent experiments, or are representative of two (C, E) independent experiments. Error bars represent SD from three (D, E, G), four (A-C), or five (F) technical replicates, and *p<0.05., **p<0.01: compared to WT BMM + WT bacteria (A, B, F, G), ^#^p<0.05, ^##^p<0.01, ^####^p<0.0001: compared to WT BMM + mutant bacteria (F, G). p-values determined by unpaired t-test.

The inability of Gal-3, -8, and -9 to restrict *Mtb* growth led us to examine whether other intracellular bacterial pathogens are restricted by the membrane damage-sensing galectins. We performed infections with *Stm, Lm,* or *Legionella pneumophila* (*Lp*), and found that TKO macrophages also exhibited normal restriction of bacterial growth with these pathogens (Fig 3E-G). TKO macrophages additionally showed no defects in IFN-γ-dependent restriction of *Stm* (Fig 3E). Furthermore, negligible differences were found between WT and TKO macrophages infected with Δ*flaA Lp* (Fig 3F), which does not activate the NAIP5-NLRC4 inflammasome [35], or an autophagy-sensitive *Lm* strain (Δ*actA* PlcA^H86A^ PlcB^H69G^, Fig 3G) [36], indicating that Gal-3, -8, and -9 are dispensable for inflammasome-independent immunity to *Lp* and *Lm*-targeted autophagy. Thus, across an array of conditions in both resting and activated macrophages, we found that Gal-3, -8, and -9 are not required for restricting the growth of several intracellular bacterial pathogens.

### TKO mice exhibit partial impairment in host resistance to chronic *Mtb* infection

To assess the role of Gal-3, -8, and -9 during *Mtb* in vivo infection, TKO and WT mice were infected with aerosolized *Mtb* (Fig 4, S4). During the initial 21 days of infection, when innate immunity predominates, there were no differences in bacterial burden in TKO mice compared to WT (Fig 4A, 4B). During chronic infection, at 9 weeks post-infection, there was a slight 2-3-fold increase in bacterial burden in the lungs of TKO mice in two independent experiments (Fig S4). Although this effect did not meet a p = 0.05 threshold for statistical significance for each experiment, pooling the CFU-fold change data from both experiments showed a statistically significant effect (Fig 4A). TKO spleens also exhibited a trend towards higher bacterial burden at this time point (Fig 4B). In addition, TKO mice succumbed slightly faster to infection than WT mice (Fig 4C). To evaluate whether these changes were dependent on inoculum size, we infected mice with a higher bacterial load (∼450 CFUs) and observed a similar trend toward increased bacterial burdens in the lungs and spleens of TKO mice (Fig S5). Taken together, these data show that mice deficient for Gal-3, -8, and -9 have no discernable defect in immunity to *Mtb* during the early acute phase of infection and have only modest defects in host resistance to chronic *Mtb* infection.

**Fig 4.**
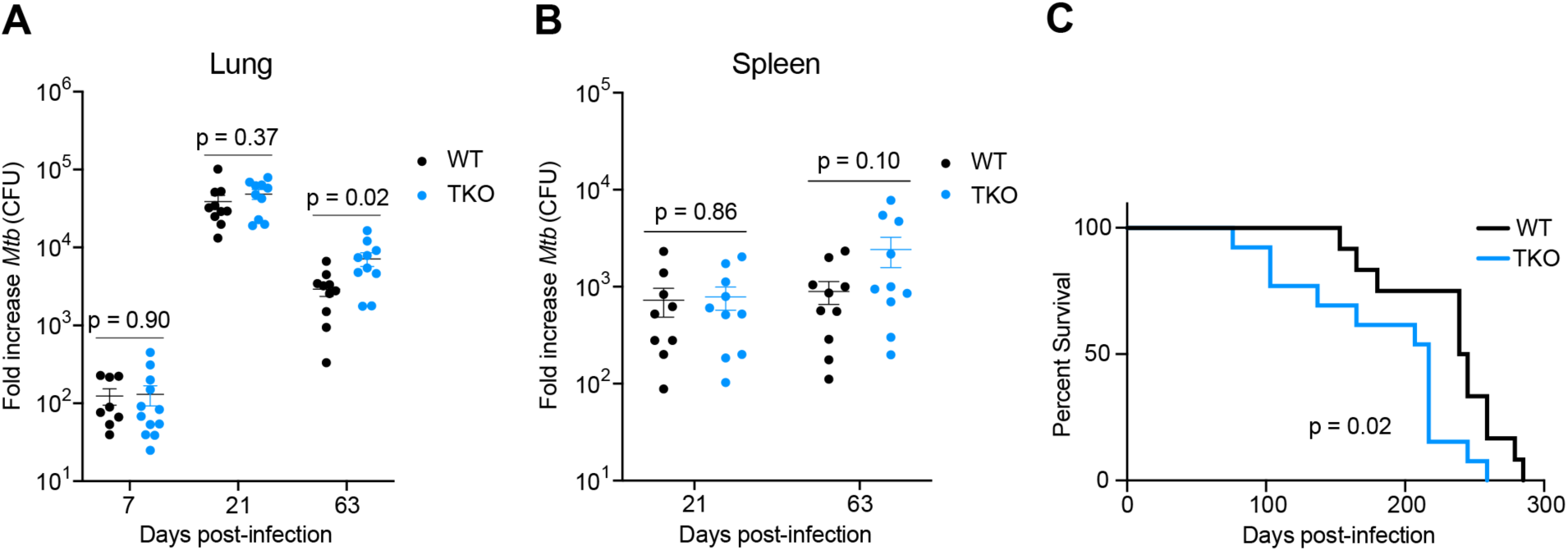
TKO mice exhibit modestly impaired resistance to chronic *Mtb* infection. (**A**) WT and TKO female mice were aerosol infected with ∼75 CFUs of *Mtb* Erdman and bacterial loads in lungs were enumerated by plating for CFU at indicated time points. CFU-fold change (relative to day 0 inoculum) from two pooled experiments is shown. *n* = 10-12 mice per genotype. (**B**) Same as in (A) but bacterial loads in spleens. (**C**) Survival of aerosol-infected mice. *n* = 12-13 mice per genotype. Figures represent one experiment (C) or represent data from two independent experiments (A, B). Bars in (A) and (B) represent the mean, error bars in (A) and (B) represent SEM, and p-values were determined by unpaired t-test (A, B), or Mantel-Cox log (C).

### Deficiency in Gal-3, -8, and -9 leads to defective antigen-specific T cell responses in the spleen

Because TKO mice exhibited resistance defects exclusively during chronic infection, when T cell-mediated immunity promotes bacterial control, we assessed whether Gal-3, -8, and -9 could play a role in shaping the adaptive immune response. To this end, we performed IFN-γ enzyme-linked immunospot (ELISpot) assays to enumerate *Mtb*-specific T cells in both WT and TKO mice infected with *Mtb.* Splenocytes were isolated 21 days post-infection when bacterial CFU in WT and TKO animals were equivalent, and ELISpot was performed with the immunodominant I-A^b^ epitopes from *Mtb* antigens ESAT-6 and Ag85B (Fig S6). Consistent with a contribution to adaptive immunity, we detected a reduction in IFN-γ-secreting Ag85B-specific T cells in TKO spleens harvested three weeks post-infection (Fig S6A). This difference seemed selective, as there were no differences in the number of IFN-γ-secreting ESAT-6-specific T cells (Fig S6A). We also measured T cell responses in the lung at this time point but found no differences between WT and TKO mice for either antigen (Fig S6B), possibly reflecting the robust enrichment of highly activated T cells in this organ [37, 38]. Finally, we measured splenic T cell responses in a high-dose infection and, in contrast to our low-dose infection results, observed no differences in Ag85B-specific T cell responses (Fig S6C), suggesting that *Mtb* infection can promote robust T cell activation in the absence of Gal-3, -8, and -9 if sufficient antigen is present. These findings suggest that membrane damage-sensing galectins promote full activation of CD4 T cells in some physiologic contexts, but not universally.

### TKO mice exhibit normal resistance to infection with *Lm* or *Stm*

Given the surprisingly mild immune defects in response to *Mtb*, we broadened our analysis and challenged WT and TKO mice with *Lm* or *Stm.* Using a *Lm* intravenous infection model, we detected similar bacterial burdens in spleen and liver from WT and TKO mice at 48 hours post-infection (Fig 5A, 5B). Likewise, bacterial burdens were unaltered in TKO mice infected with *Stm* via the intraperitoneal route (Fig 5C, 5D). Thus, despite their well-established role in membrane damage-sensing and our data demonstrating direct pathogen recognition, simultaneous loss of all three phagosome-localized galectins has no discernable impact on the ability of the host to restrict several bacterial pathogens during the acute phase of infection and results in only modest impairments to host immunity during chronic *Mtb* infection.

**Fig 5.**
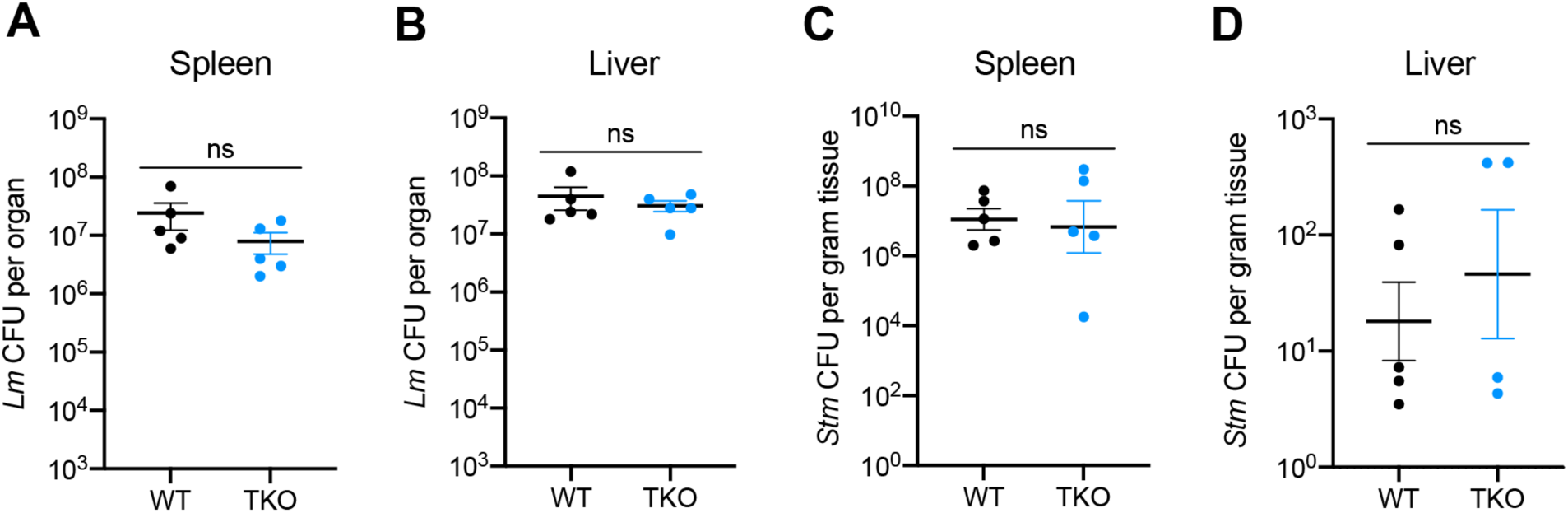
TKO mice exhibit normal bacterial control during infection with *Lm* or *Stm*. (**A**) WT and TKO mice were intravenously infected with 1 x 10^5^ CFUs of *Lm* and bacterial loads in spleens were enumerated by plating for CFU at 48 hours post-infection. *n* = 5 mice per genotype. (**B**) Same as in (A) but bacterial loads in livers. (**C**) WT and TKO mice were infected via the intraperitoneal route with 1000 CFUs of *Stm* and bacterial loads in spleens were enumerated by plating for CFU at 24 hours post-infection. *n* = 5 mice per genotype. (**D**) Same as in (C) but bacterial loads in livers. *n* = 4-5 mice per genotype. Figures represent data from one experiment (A, B), or are representative of three independent experiments (C, D). Bars represent the mean, error bars represent SEM, and p-values were determined by unpaired t-test.

## Discussion

Gal-3, -8, and -9 have been implicated in pathogen sensing, autophagy activation, and membrane repair [9,15–18,20]. Our results showed that TKO macrophages exhibit a reduction in *Mtb* delivery to the lysosome and increased *Mtb* colocalization with p62 and ubiquitin. While these data suggest that galectin deficiency leads to a partial block in the autophagy pathway and consequently an accumulation of p62- and ubiquitin-associated bacteria, it is possible that defects in phagosome repair also expose bacteria to increased p62 and ubiquitin targeting. Robust ubiquitin recruitment to *Mtb* in TKO macrophages also suggests that ubiquitin-mediated autophagy pathways, promoted via Parkin, Smurf1, Trim16, or cGAS/STING [15,30,31,33], are largely intact in TKO macrophages and may compensate for defects in galectin-driven autophagy to promote bacterial growth restriction.

Previous reports demonstrated that Gal-3 and Gal-8 can promote restriction of bacterial growth in immortalized cell lines [9,15,20]. We were surprised that in our experiments, TKO macrophages had no discernable defects in their ability to restrict growth of *Stm, Lm, Lp,* or *Mtb* in either the resting state or following IFN-γ activation. Taken together, this suggests that while membrane damage-sensing galectins have the potential to restrict bacterial growth in cell lines, there may be key differences between these cells and primary macrophages that account for the differential requirements for galectin activity to restrict bacterial growth.

Given the previous work with individual galectins during intracellular bacterial infection, we were surprised that mice lacking Gal-3, -8, and -9 had no resistance defects during the acute stages of infection with *Stm, Lm,* and *Mtb* and only a modest susceptibility defect during the chronic stage of *Mtb* infection. There are several possible explanations for the seemingly minimal contribution of Gal-3, -8, and -9 to antibacterial immunity during acute infection. First, redundancy in galectin function (e.g. different galectins or other pathways) may be compensating for Gal-3, -8, and -9 deficiency, although no other galectins are currently known to be targeted to damaged membranes.

Alternatively, as several intracellular pathogens actively inhibit autophagy using specific virulence factors [39–41], these pathogens may inactivate galectins or downstream autophagy steps as a virulence strategy. Finally, it is possible that galectins play roles in cell types other than macrophages or in different physiologic contexts, such as mucosal immunity in the gut, where epithelial cells rather than professional phagocytes are the initial line of defense against an array of pathogens. Autophagy plays a protective role in the gut epithelium [42], and certain innate immune mechanisms are only evident during infection of the gut epithelium, exemplified by NOD2-mediated restriction of *Lm* growth during oral but not intravenous infection [43]. Thus, galectins may still have an important antibacterial role in other physiologic contexts beyond those we have tested here.

We did observe modest defects in bacterial control and disease susceptibility during the chronic stage of *Mtb* infection in TKO mice. This chronic phase phenotype is similar to the autophagy-related phenotype in *Mtb-*infected *Smurf1-/-* mice, which also have higher *Mtb* burdens exclusively during the chronic phase and exhibit a very similar survival phenotype [31]. It is possible that TKO mice exhibit defects in resistance due to aberrant inflammatory responses previously reported in autophagy mutant mice [31,44,45]. Together, these findings suggest a role for certain autophagy-related events during chronic tuberculosis infection when there is partial containment of *Mtb* by coordinated innate and adaptive immune responses.

Our finding that galectin deficiency leads to slight impairments in both autophagy and antigen-specific CD4 T cell activation raises the possibility that these phenotypes could be linked and suggests how an autophagy phenotype might manifest during later stages of infection. Autophagy mediates the capture and degradation of cytosolic pathogens and antigens, after which the peptide constituents are trafficked to MHC-class-II-containing late endosomes and then loaded onto MHC class II complexes for presentation to CD4 T cells [46–49]. During *Mtb* infection, inhibition of autophagy was shown to impair Ag85B-specific, but not ESAT-6-specific CD4 T cell activation [50], possibly suggesting that some microbial antigens, but not others, require autophagy for efficient antigen presentation. Similarly, the CD4 T cell activation defect we observed in TKO mice was specific to Ag85B, while ESAT-6-specific CD4 T cell responses were unchanged. How some antigens, but not others, might access MHC class II compartments via autophagy remains unknown. It is possible that highly expressed and secreted antigens, like ESAT-6 [51], do not require autophagy for antigen presentation and instead access the antigen presentation pathway via phagocytosis and endolysosomal trafficking. In contrast, Ag85B, a cell wall antigen that is downregulated at 3 weeks post-infection and exhibits low abundance during chronic infection [52, 53], might rely on autophagy for efficient antigen presentation.

Although we and Wu et al. found that Gal-9 can bind directly to the surface of *Mtb* [25], the ligands recognized by Gal-9 during infection remain unclear. The mycobacterial cell wall, from inside to out, is comprised of an inner membrane, a layer of peptidoglycan covalently linked to a subsequent layer of arabinogalactan (AG), an extremely hydrophobic and relatively-impermeant outer membrane (OM) consisting mainly of long-chain mycolic acids, and a loosely associated polysaccharide “capsule” at the very surface [54, 55]. Wu et al. reported that Gal-9 recognized *Mtb* AG [25], though this layer is internal to the OM and unlikely to be accessible to Gal-9 in healthy, dividing bacteria [55]. A possible solution to this conundrum is that permeabilization of the outermost capsule and mycolic acid layer in vivo (or with detergents routinely used in mycobacterial growth medium) [54], might expose AG to Gal-9. Moreover, it is possible that Gal-9 primarily recognizes dead or dying bacteria that have already been damaged during infection in order to clear bacterial corpses (and their PRR-activating ligands) from the cytosol via autophagy. Gal-9 localizes to damaged endosomes and lysosomes independently of bacterial infection [9, 18], indicating that Gal-9 also recognizes an endogenous carbohydrate ligand. In line with this, our immuno-EM studies showed that most Gal-9 colocalizes with host membranes, while a small fraction colocalizes with *Mtb* in infected macrophages. Recognition of host glycoproteins on damaged membranes or release of AG or other Gal-9 bacterial ligands could localize Gal-9 to extra-phagosomal compartments [56]. Lastly, our observation that Gal-9 also binds *Stm* but not *Lm* suggests Gal-9 recognition of bacterial constituents is not limited to AG and is selective for a subset of bacteria. Indeed, other galectin-bacteria interactions have been identified, including Gal-3/Gal-9-*Leishmania major*, Gal-4/Gal-8-*Escherichia coli,* and Gal-3-*Neisseria meningitidis* [22–24]. This suggests a scenario where galectins have divergent binding preferences, allowing the recognition of a broader array of microbes.

In conclusion, our data suggest that although Gal-3, -8, and -9 contribute to autophagy in macrophages, they play a limited role in the immune response to systemic bacterial infection. These membrane damage-sensing galectins had no discernable impact on acute infection with *Mtb*, *Stm*, or *Lm* in vivo, and had a modest effect on host resistance to chronic *Mtb* infection. These data suggest that the role of Gal-3, -8, and -9 in antibacterial immunity is not generalized, but rather is limited to specific physiologic contexts in vivo.

## Methods

### Ethics Statement

Mouse use and procedures were approved by the Office of Laboratory and Animal Care at the University of California, Berkeley (protocol AUP-2015-11-8096), and University of California, Davis (protocol #22365) in adherence with federal guidelines from the *Guide for the Care and Use of Laboratory Animals* of the National Institute of Health.

### Pathogen pull-down for mass spectrometry and immunoblotting

For *Mtb* pull-down mass spectrometry, *Mtb* was fixed in PBS with 4% PFA, washed three times with PBS, and pelleted. 50 μL of *Mtb* pellet was incubated for 3 hours with 1 mL of 10 mg/mL differentiated THP-1 lysate, washed four times with PBS, and eluted in 8M urea with Rapigest (Waters) before LC-MS sample preparation. For *Mtb*, *Lm, Stm*, and *Cn* pull-down for Gal-9-FLAG immunoblotting, pathogens were fixed, washed, and pelleted as described above. 1 mL of 10 mg/mL RAW 264.7 Gal-9-FLAG lysate was incubated, washed, and eluted as described above prior to immunoblotting.

### Mass spectrometry

Samples were denatured and reduced in 2 M urea, 10 mM NH4HCO3, 2 mM DTT for 30 minutes at 60°C, then alkylated with 2 mM iodoacetamide for 45 minutes at room temperature. Trypsin (Promega) was added at a 1:100 enzyme:substrate ratio and digested overnight at 37°C. Following digestion, samples were concentrated using C18 ZipTips (Millipore) according to the manufacturer’s specifications. Desalted samples were evaporated to dryness and resuspended in 0.1% formic acid for mass spectrometry analysis.

Peptides were resuspended in 0.1% formic acid and 3% directly injected on a 75 μm ID column (New Objective) packed with 25 cm of Reprosil C18 3 μm, 120 Å particles (Dr. Maisch). Peptides were eluted in positive ion mode into an Orbitrap Elite mass spectrometer (Thermo Fisher) via a Nanospray Flex Ion Source (Thermo Fisher). Elution of peptides was achieved by an acetonitrile gradient delivered at a flow rate of 400 nL/min by an Easy1000 nLC system (Thermo Fisher). All mobile phases contained 0.1% formic acid as buffer A. The total gradient time was 70 minutes, during which mobile phase buffer B (0.1% formic acid in 90% acetonitrile) was ramped from 5% to 30% B over 55 minutes, followed by a ramp to 100% buffer B over a subsequent 5 minutes, and held at 100% for 10 minutes to wash the column. MS data was collected over the first 60 minutes of the gradient. The ion transfer tube was set to 180°C and the spray voltage was 1500V. MS1 scan was collected in the ion trap in centroid mode with 120K resolution over a scan range of 200-2000 m/z, an S-Lens RF of 68%, a 50 ms maximum injection time, and a AGC target of 3e4. Species with a charge state z=1 or unassigned charge states were excluded from selection for MS/MS. Resonance excitation MS/MS was performed on the 20 most abundance precursors with a minimum signal intensity of 200, an isolation width of 2 m/z, a 0.25 activation Q, a 10 ms activation time, a 100 ms maximum injection time, and an AGC target of 1e3. Dynamic exclusion was employed to exclude a list of 50 previously selected precursors within 180 seconds and using a +/-1.5 Da window. The results raw data was matched to protein sequences as previously described [57].

### Western Blot

Cells were lysed using RIPA buffer (Pierce) supplemented with Complete Mini EDTA-free Protease Inhibitor (Roche). Lysate protein concentration was determined using BCA protein assay kit (Pierce). Proteins were separated with denaturing PAGE 4-20% Mini-PROTEAN TGX Precast Protein gel (Bio-Rad) and transferred to Trans-Blot Turbo Mini Nitrocellulose (Bio-Rad) using Trans-Blot Turbo Transfer System (Bio-Rad). Membranes were blocked with LI-COR Odyssey blocking buffer for 1 hour at room temperature with agitation. Primary antibodies were added and incubated overnight at 4°C with agitation. Primary antibodies used were anti-Gal-3 (SCBT sc-32790, primary 1:1000, secondary 1:5000), anti-Gal-8 (Abcam ab69631, primary 1:500, secondary 1:5000), anti-Gal-9 (Abcam ab69630, primary 1:500, secondary 1:5000), anti-β-actin (SCBT sc-47778, primary 1:1000, secondary 1:5000), anti-FLAG (Sigma F3165, primary 1:1000, secondary 1:5000). Secondary antibodies used were LI-COR IRDye 680RD or 800CW goat anti-mouse IgG or anti-rabbit IgG and used at dilutions indicated above. Following primary stain, blots were washed three times 5 minutes each in PBS or PBS 0.1% Tween 20 (PBS-T). Secondary antibodies were incubated for 30 minutes at room temperature with agitation. Blots were washed three times prior to imaging on a LI-COR Odyssey imaging system. PBS-T was used for washing all immunoblots, except anti-Gal-8 and anti-Gal-9 immunoblots, which used PBS no tween for washes. Blots were imaged on a LI-COR Odyssey imaging system.

### Bacterial and fungal cultures

#### Mycobacterium tuberculosis

All *Mtb* experiments were performed with *Mtb* Erdman strain or strains derived from *Mtb* Erdman strain. Low passage frozen stocks of *Mtb* were grown to log phase in 7H9 (BD) liquid media supplemented with 10% Middlebrook OADC (Sigma), 0.05% Tween-80 and 0.5% glycerol in roller bottles at 37°C. *Mtb-*LUX expressing luciferase from the *luxCDABE* operon has been described previously and was cultured as described above [34]. WT or Δ*eccC Mtb* expressing eGFP under control of the MOP promoter was a gift from Dr. Sarah Stanley’s laboratory [58], and was cultured as described above. For BMM infections, log phase bacteria were washed twice with PBS and sonicated three times at 90% amplitude for 30 seconds each then diluted into BMM media (DMEM with 10% FBS, 2 mM L-glutamine, 10% MCSF, 11 mg/mL sodium pyruvate) prior to infection. For THP-1 and RAW 264.7 infections, 10 mL of the *Mtb* culture was washed twice in PBS with 10% heat-inactivated horse serum. After washing, *Mtb* was resuspended in 5 mL PBS with 10% heat-inactivated horse serum and sonicated twice at 50% amplitude for 15 seconds. *Mtb* was diluted into DMEM with 10% heat-inactivated horse serum, and this inoculum was added to THP-1 cells prior to spinfection.

### Salmonella enterica serovar Typhimurium

For pathogen pull-down experiments, *Stm* strain SL1344 was cultured overnight at 37°C with shaking in Luria-Bertani (LB) broth and backdiluted 1:100 1 day prior to infection, then allowed to recover to mid-log. BMM infections were performed with *Stm* strain IR715. For mouse infections, *Stm* strain IR715 was cultured in LB broth overnight at 37°C with shaking.

#### Listeria monocytogenes

All *Lm* strains used in this study were derived from strain 10403S (streptomycin-resistant). Δ*actA* PlcA^H86A^ PlcB^H69G^ has been previously described [36]. For macrophage infections, *Lm* was grown to stationary phase overnight, slanted at 30°C in filter-sterilized BHI media with 200 μg/mL streptomycin then diluted into PBS for OD600 measurement and inoculum preparation.

#### Legionella pneumophila

All *Lp* strains used are JR32 and express the *Photorhabdus luminescens luxCDABE* operon. *LuxCDABE*-expressing WT or isogenic mutant for *flaA* have been previously described [59]. Bacteria were cultured in Charcoal-Yeast Extract Agar (CYE, 10 g/L 4-morpholinepropanesulfonic acid [MOPS], 10 g/L Yeast extract, 15 g/L technical agar, 2 g/L activated charcoal, supplemented with 0.4 g/L L-cysteine and 0.135 g/L Fe(NO3)3) at 35-37°C for 4 days from frozen stocks. Single colonies were grown on fresh plates for another 2 days. For macrophage infections, bacteria were grown on solid plates and resuspended in autoclaved water and diluted in RPMI.

#### Cryptococcus neoformans

*Cn* Serotype A strain H99 was grown in yeast culture conditions on YPAD medium [1% yeast extract, 2% Bacto-peptone, 2% glucose, 0.015% L-tryptophan, 0.004% adenine] at 30°C. Cells were harvested mid-log phase for in vitro binding studies.

### Cell culture

RAW 264.7 macrophages were cultured in DMEM, 10% FBS, 2 mM L-glutamine, 10 mM HEPES. RAW 264.7 macrophages were transduced with lentiviral particles containing Gal-9-3xFLAG expressed from pLenti-puro (Addgene #39481). Primary murine bone marrow-derived macrophages (BMMs) were isolated and cultured in BMM media (DMEM with 10% FBS, 2 mM L-glutamine, 10% MCSF, 11 mg/mL sodium pyruvate) as previously described [26]. For all experiments, BMMs were thawed and allowed to equilibrate for three days prior to replating. THP-1 cells were cultured in RPMI 1640 medium supplemented with 1 mM sodium pyruvate, 10mM HEPES, 4.5 g/L glucose, 0.05 mM 2-mercaptoethanol, 2 mM L-glutamine, 5% fetal calf serum, and 5% bovine growth serum. THP-1 cells were maintained at a concentration between 2e5 cells/mL and 1e6 cells/mL.

### LLOMe treatment

BMMs were incubated with 50 nM Lysotracker Red DND-99 (Thermo Fisher) for 3 hours prior to incubation with LLOMe (Cayman Chemical) at 0.5 mM for either 3 hours, or 30 minutes followed by 30 minute washout. Cells were fixed for 15 minutes in PBS with 4% PFA followed by three PBS washes for 5 minutes each. Cells were then DAPI stained for 30 minutes and imaged.

### THP-1 shRNA generation

shRNAs were designed to target human Galectin-9 mRNA and cloned into the pLKO.1 vector (Addgene #10878). The sense shRNA sequences are CCTGGTGCAGAGCTCAGATTT (shGal-9 #1) and GCTCTGTGCAGCTGTCCTA (shGal-9 #2). Lentiviral particles were produced in Lenti-X cells (Clontech) per manufacturer’s instructions. Early passage THP-1 cells were transduced with lentiviral particles containing the shRNA expression vector. The cells were selected in 0.5 μg/mL puromycin to obtain a polyclonal population. Knockdown efficiency was assessed via qPCR using the following primers: 5’-CCGAAAAATGCCCTTCGTCC-3’ (forward), 5’-ACCTTGAGGCAGTGAGCTTC-3’ (reverse) and determined to be 95% and 91% knockdown for shGal-9 #1 and shGal-9 #2, respectively.

### Macrophage infections

#### Mycobacterium tuberculosis

To ensure bacterial replication or autophagy targeting was not impacted by differences in macrophage concentration, WT BMMs were plated in 96-well plates at the target cell density, while TKO BMMs were plated on a cell density gradient in order to ensure equivalent plating numbers of each genotype. After plating, cells were allowed to adhere and equilibrate for 2 to 3 days prior to infection. On day 0 of infection, one plate was fixed in PBS with 4% PFA, washed, DAPI stained, and nuclei were counted at 5X magnification on an Opera Phenix microscope, thus allowing determination of cell density-matched wells. During analysis, wells of different genotypes were only compared if cell counts were matched within <5% of each other. For microscopy, CFU, and *Mtb-*LUX experiments, cells were infected by removing media from wells, overlaying cell monolayers with bacterial suspensions, and spinfecting for 10 minutes at 300 rcf.

For *Mtb*-LUX infections, cells were plated at 4 x 10^4^ cells per well (target density) in Nunc 96-well white walled TC treated plates (Thermo Fisher). For IFN-γ treatment, cells were incubated with 1.5 ng/mL IFN-γ the day before infection. BMMs were infected at a MOI of 1 or 5. Luminescence of *Mtb*-LUX-infected BMMs in cell density-matched wells was measured as previously described [57]. For all *Mtb* microscopy experiments, WT *Mtb*-GFP or Δ*eccC Mtb*-GFP were used. BMMs were plated in either 24-well glass-bottom plates (Cellvis) at 2 x 10^5^ cells per well, or 96-well PerkinElmer Cell Carrier Ultra plates at 2.5 x 10^4^ cells per well (target density). All infections for microscopy used an MOI of 2. At indicated time points, microscopy plates were washed in warm PBS, fixed in PBS with 4% PFA, washed three times with PBS, and stored in PBS at 4°C before immunostaining. For lysotracker experiments, cells were stained with 50 nM Lysotracker Red DND-99 in BMM media for 3 hours prior to fixation and imaged immediately after washing out the fixative.

For *Mtb* CFU experiments, BMMs were plated in 24-well TC treated plates at 2 x 10^5^ cells per well, infected with WT *Mtb* Erdman strain using an MOI of 2, and *Mtb* CFU enumeration was performed as previously described [58].

For *Mtb* infection of THP-1 cells, THP-1 cells were differentiated for 3 days in DMEM with 2 mM L-glutamine, 1 mM sodium pyruvate, and 10% heat-inactivated fetal calf serum with 10 ng/mL PMA and 0.1 ng/mL Vitamin D. On day 0 of differentiation, THP-1 cells were plated at 4 x 10^4^ cells per well in a 96-well plate. The media was exchanged with fresh differentiation media daily. Bacterial suspension was prepared as described above and THP-1 cells were spinfected for 5 minutes. Cells were incubated at 37°C for 30 minutes. THP-1 cells were washed 3 times with PBS with 1% heat-inactivated horse serum. Differentiation media lacking phenol red with PMA and Vitamin D was replaced after the washes. Luminescence of *Mtb*-LUX-infected THP-1 cells was measured as previously described [57]. Half of the media was discarded and replaced with fresh differentiation media with PMA and Vitamin D on days 1, 3, and 5 post-infection.

### Salmonella enterica serovar Typhimurium

1 x 10^5^ BMMs were plated on 24-well plates 24 hours prior to infection. For IFN-γ treatment, cells were incubated with 0.01 μg/mL IFN-γ the night before infection. Bacteria grown to mid-log phase as described above were pelleted and resuspended in BMM media. Suspension was incubated at 37°C for 30 minutes for opsonization. Bacteria were then pelleted and resuspended to 1 x 10^9^ CFU/mL prior to spinfection with BMMs for 5 minutes at 250 rcf followed by 37°C incubation for 30 minutes for phagocytosis. Cells were washed with PBS and incubated with BMM media with 100 μg/mL gentamicin at 37°C incubation for 30 minutes, then washed with PBS and incubated with BMM media with 5 μg/mL gentamicin. For CFU enumeration, cells were washed with PBS and lysed in 500 uL water for 10 minutes prior to lysate serial dilution and plating on LB agar plates.

#### Listeria monocytogenes

A total of 3 x 10^6^ BMMs were plated on 14 coverslips in 60 mm non-TC treated dishes and infected the next day at an MOI of 0.25. *Lm* CFU enumeration was performed as previously described [60].

#### Legionella pneumophila

Bacterial replication was measured using a luminescence-based replication assay as previously described [61]. Briefly, 10^5^ WT or TKO BMMs were plated on 96-well plates and incubated overnight. Medium was replaced with bacterial suspension in BMM media at an MOI of 0.01. At indicated time points, luminescence emission was measured at 470 nm with a Spectra-L plate reader (Molecular Devices).

### Immunofluorescence

Following fixation, washing, and storage in PBS at 4°C, cells were blocked and permeabilized in PBS with 5% FBS or 2% BSA and 0.1% Triton X-100 for 30 minutes at room temperature. Cells were then stained in blocking buffer overnight at 4°C with the following primary antibodies at the indicated dilutions: anti-Gal-9 (Abcam ab275877, primary 1:100, secondary 1:500), anti-Gal-3 (SCBT sc-32790, primary 1:50, secondary 1:500), anti-polyubiquitin (Millipore Sigma ST1200, primary 1:400, secondary 1:2000), anti-ubiquitin K63-specific (Millipore Sigma 05-1308, primary 1:200, secondary 1:2000), anti-ubiquitin K48-specific (Millipore Sigma 05-1307, primary 1:200, secondary 1:2000), anti-p62 (Abcam ab109012, primary 1:400, secondary 1:4000), anti-TAX1BP1 (Bethyl Laboratories A303-791A, primary 1:200, secondary 1:1000), and anti-OPTN (Bethyl Laboratories A301-829A, primary 1:200, secondary 1:1000). Cells were then washed three times with PBS for 5 minutes each. DAPI and secondary antibody staining was performed at room temperature for 30 minutes using anti-mouse or anti-rabbit Alexa Fluor 647 (Invitrogen) at the above indicated dilutions. Cells were washed three times with PBS for 5 minutes then stored at 4°C or imaged.

### Automated confocal microscopy and image analysis

All microscopy experiments were performed with an Opera Phenix High-Content Screening System confocal microscope (PerkinElmer). Microscopy plates were imaged at 10X magnification for nuclei counting and cell density matching, 40X or 63X magnification for colocalization or lysotracker puncta analysis, or 63X magnification for representative images. Automated analysis for colocalization, nuclei counting, and lysotracker puncta experiments was performed with PerkinElmer Harmony software package. An analysis script was created to measure colocalization of *Mtb* and indicated markers from >4000 bacteria.

### Electron-microscopy

RAW 264.7 cells expressing mouse Gal-9-3xFLAG were infected with *Mtb* at an MOI of 1 for 6 hours. Cells were then detached using 1 mM EDTA in PBS and fixed for 10 minutes at room temperature using freshly prepared periodate-lysine-paraformaldehyde stain (0.2 M lysine-HCl pH 7.4, 4% paraformaldehyde, 0.1 M sodium m-periodate). Following fixation, cells were pelleted and resuspended in PBS without washing and stored at 4° until embedding was performed. Samples were embedded in 10% gelatin and infiltrated overnight with 2.3 M sucrose/20% polyvinyl pyrrolidone in PIPES/MgCl_2_ at 4°C. Samples were trimmed, frozen in liquid nitrogen, and sectioned with a Leica Ultracut UCT7 cryo-ultramicrotome (Leica Microsystems). Ultrathin sections of 50 nm were blocked with 5% FBS/5% NGS for 30 minutes and subsequently incubated with mouse anti-FLAG (Sigma F3165) for 1 hour at room temperature. Following washes in block buffer, sections were incubated with goat anti-mouse IgG + IgM 18 nm colloidal gold conjugated secondary antibody (Jackson ImmunoResearch Laboratories) for 1 hour. Sections were stained with 0.3% uranyl acetate/2% methyl cellulose and viewed on a JEOL 1200 EX transmission electron microscope (JEOL USA) equipped with an AMT 8 megapixel digital camera and AMT Image Capture Engine V602 software (Advanced Microscopy Techniques). All labeling experiments were conducted in parallel with controls omitting the primary antibody.

### Mice

All mice used were specific pathogen free, maintained in 12 hour light-dark cycle, and given standard chow diet ad libitum. 8 to 12-week-old male and female mice were used. WT C57BL/6J mice were obtained from Jackson Laboratories (JAX). CRISPR/Cas9 targeting was performed by electroporation of Gal-3, Gal-8, and Gal-9-targeting Cas9-sgRNA RNP complexes into fertilized zygotes from C57BL/6J female mice (JAX, stock no. 000664) [62]. TKO mice were generated by targeting exon 1 of each galectin using two guides per exon; sgRNA sequences: 5’-GGCTGGTTCCCCCATGCACC-3’ and 5’-CTCCAGGGGCAGTTGGGCCG-3’ (Gal-3), 5’-CAGCTAGACCTTTTGAACCG-3’ and 5’-CACCATGAACACGATCTCAA-3’ (Gal-8), and 5’-TACCCTCCTTCCTCAAACCG-3’ and 5’-GAACGGACAGTGGGGTCCTG-3’ (Gal-9). Founder mice were genotyped by PCR amplifying the targeted exons from mouse tail DNA, followed by Sanger sequencing to identify mice carrying mutations. The following primers were used: Gal-3 PCR fwd tgtgaatcttctcccatgtcccagc; Gal-3 PCR rev tccttcttaccagtggtccagc; Gal-3 seq agatcacaaatgcctgtagtc; Gal-8 PCR fwd tccatataagccagctcatgctgtgg; Gal-8 PCR rev gcgctcctagaagaagtaagacctaagg; Gal-8 seq ctaaggtttatctgttccatctgg; Gal-9 PCR fwd atgacattgccttccacttcaacc; Gal-9 PCR rev tcaaagggcatccccttctgg; Gal-9 seq cacctcaggcagtcag. Mice with mutations were bred to WT C57BL/6J mice to separate modified haplotypes. Homozygous lines were generated by interbreeding heterozygotes carrying matched haplotypes. Two independent TKO mouse lines were established. TKO strain 1 (Gal-3, 34nt deletion; Gal-8, 1nt insertion; Gal-9, 58 nt deletion) was used for the *Mtb*, *Lm*, and *Lp* experiments. TKO strain 2 (Gal-3, 5nt deletion; Gal-8, 103nt deletion; Gal-9, 17nt deletion) was used for the *Stm* macrophage and mouse experiments.

### Mycobacterium tuberculosis aerosol infection

To prepare infection inoculum*, Mtb* Erdman strain frozen stock was diluted in sterile water and sonicated at 90% amplitude three times for 30 seconds. 8 to 12-week-old male and female mice were infected with approximately ∼75 CFUs (low-dose) or >400 CFUs (high-dose) of prepared inoculum using a Glas-Col inhalation exposure system. At indicated time points, lungs and spleens were isolated, homogenized, and plated on 7H10 plates with Polymyxin B (200,000 U/L), Carbenicillin (50 mg/L), Trimethoprim lactate (20 mg/L) and Amphotericin B (5 mg/L). Plates were incubated at 37°C for 25 - 28 days before counting colonies. For survival experiment, mouse weight was measured weekly and mice were sacrificed after losing 15% of maximum body weight.

### *Salmonella enterica* serovar Typhimurium intraperitoneal infection

*Stm* strain IR715 was cultured as described above. Mice received 100 μl of sterile PBS or 1000 CFUs diluted in 100 μl PBS by intraperitoneal injection. Mouse weights were monitored daily during infection. Systemic bacterial levels were characterized at day 1, 2, or 3 post-infection by harvesting liver and spleen and enumerating CFU. Liver and spleen were collected in PBS, weighed, and homogenized. Homogenates were serially diluted and plated on LB agar plates. CFU per gram tissue was calculated after overnight growth at 37°C.

### Listeria monocytogenes intravenous infection

*Lm* was grown at 37°C with shaking at 200 rpm to mid-log phase. Bacteria were washed in PBS and resuspended at 5 x 10^5^ colony-forming units (CFU) per 1 mL of sterile PBS. Mice were then injected with 1 x 10^5^ CFU via the tail vein. 48 hours post-infection, livers and spleens were collected, homogenized, and plated on LB streptomycin 200 μg/mL to determine the number of CFU per organ.

### Interferon-γ enzyme-linked immunospot (ELISpot)

Spleens were isolated and pressed through a 70 μm cell strainer to generate a single cell suspension. Lung cells were isolated and digested in RPMI with Liberase TM (70 μg/ml) and DNase I (30 μg/ml) in gentleMACS C-tubes then placed at 37°C for 30 minutes, followed by gentleMACS tissue homogenization for single cell suspension. Cells were pelleted and RBCs were lysed using ACK lysing buffer. Cells were washed, pelleted, resuspended in X-VIVO TM 15 (Lonza) and counted on a hemocytometer or using fluorescent count beads on an SH800 Sony Sorter. Cells were plated, stimulated overnight with indicated peptides (2 μg/ml) or Concanavalin A (ConA, 2.5 μg/ml), developed, and analyzed as previously described [63]. ConA-stimulated cells were used as a positive control and to determine total cell counts. IFN-γ spot-forming cell (SFC) counts from peptide-stimulated wells were normalized to SFC counts from ConA-stimulated wells to control for differences in cell density between samples.

### Statistical analysis

Analysis of statistical significance was performed with Prism 9 (GraphPad).

## Acknowledgments

We thank Erika Oki, Justin Paluba, Rita McCall, Miles Tuncel, Laura Flores, Christopher Noel, Dmitri Kotov, Danielle Swaney, Hannah Nilsson, Michael Cronce, Allison Roberts, and Chris Rae for technical assistance. We thank Dr. Mary West and Dr. Deepa Sridharan of the High-Throughput Screening Facility (HTSF) at UC Berkeley. This work was performed in part in the HTSF, which provided the Opera Phenix microscope, funded by NIH S10OD021828. The schematic was created with BioRender.com. Finally, we thank James Olzmann and members of the Cox, Vance, and Stanley labs for helpful discussions. This work was supported by NIH grants P01 AI063302 (JSC, SAS, DAP, REV), R37 AI075039 (REV), R01 AI27655 (DAP), R01 AI144149-04 (BHP). REV is an Investigator of the Howard Hughes Medical Institute.

## Author Contributions

Conceptualization: HMM, BHP, JSC

Methodology: HMM, JC, RRL, JRJ, WLB, SRM, BHP

Data Curation, Software: JRJ Analysis: HMM, JC, JRJ, BHP

Investigation: HMM, JC, RRL, CED, JRJ, GRG, EVD, WLB, SRM, TR, IS, AYL, BHP

Writing – original draft: HMM

Writing – review & editing: HMM, BHP and JSC with input from all co-authors Funding Acquisition: REV, SAS, DAP, BHP, JSC

Supervision: REV, SAS, DAP, NJK, BHP, JSC

---

**Fig S1.**
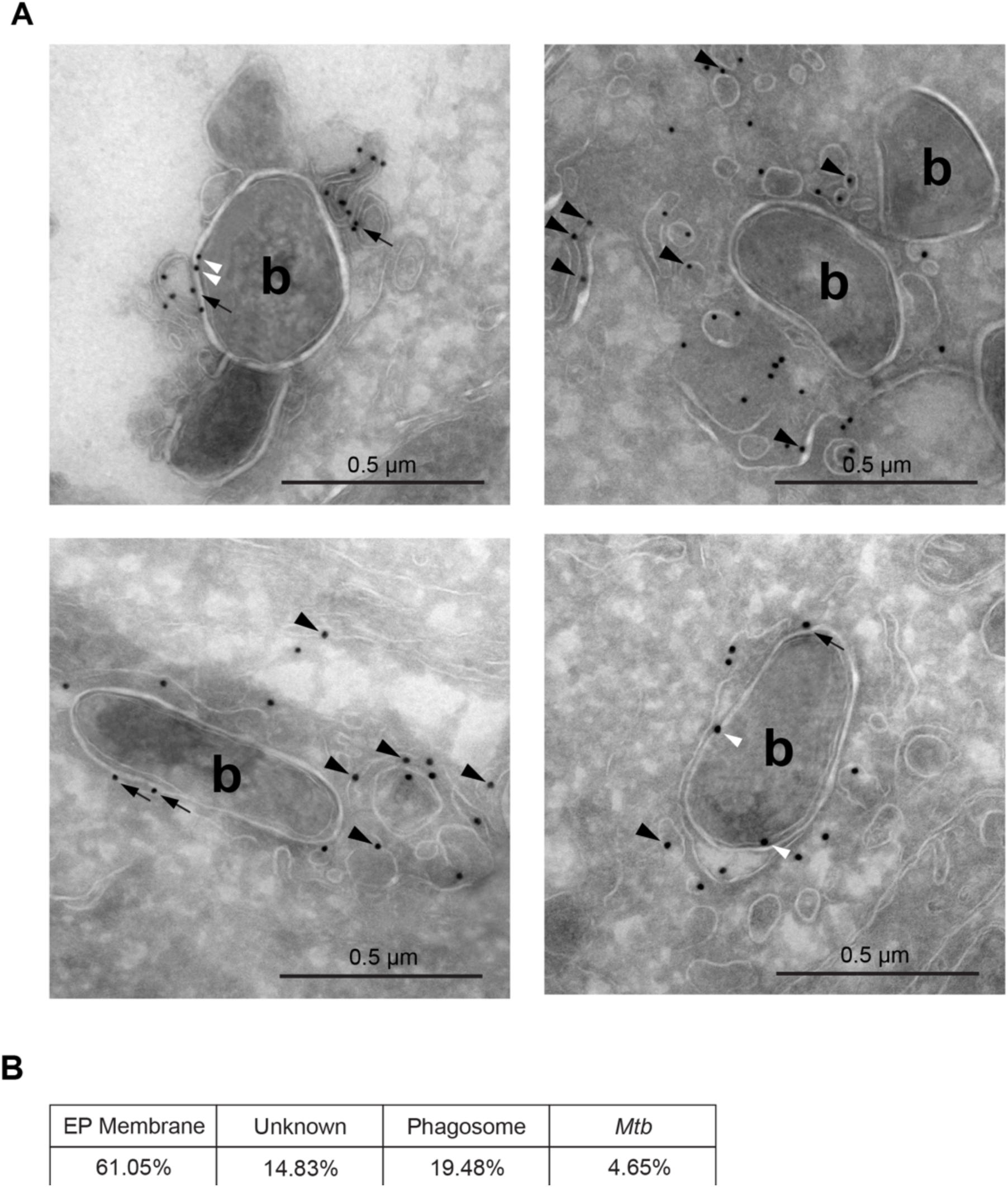
Gal-9 localization during *Mtb* infection. (**A**) Cryosections of RAW 264.7 cells stably expressing Gal-9-FLAG infected with WT *Mtb* (MOI = 1) 6 hours post-infection. Gal-9 localization to extra-phagosomal membranes (black arrowheads), phagosomes (black arrows), and *Mtb* (white arrowheads); b, bacteria. Four representative micrographs are shown from a dataset of 26 total images. Scale bar = 0.5 μm. (**B**) Quantification of (A) for Gal-9-FLAG localization to indicated structures; EP membrane, extra-phagosomal membranes. Values are a percent of total Gal-9-FLAG puncta in the dataset.

**Fig S2.**
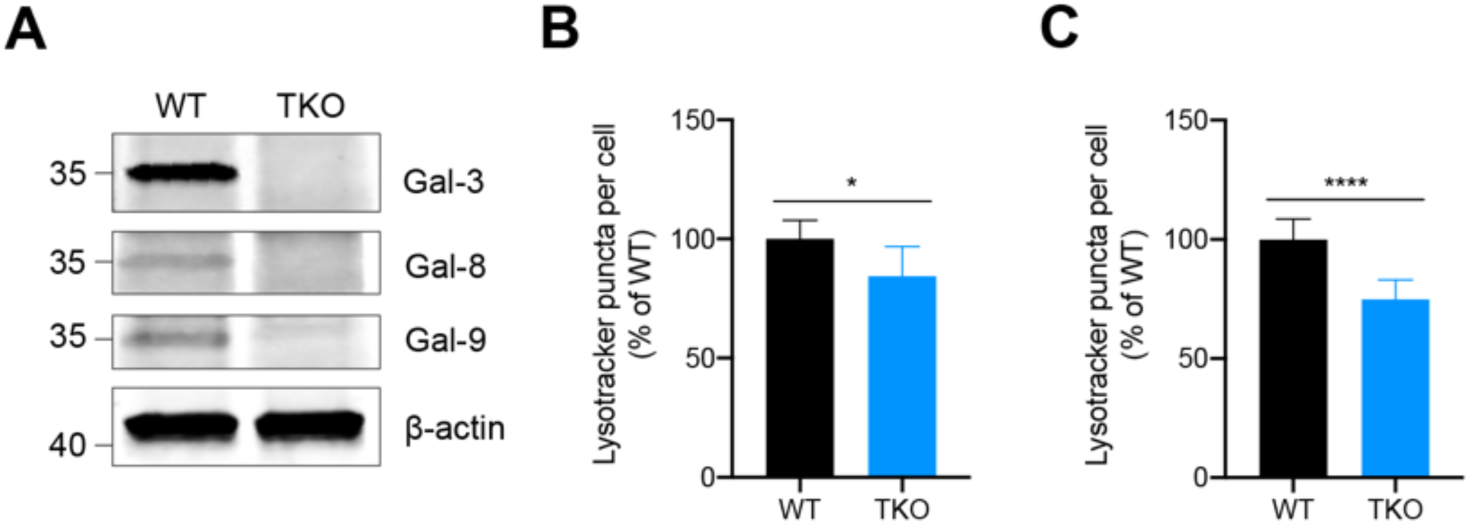
Generation and validation of Gal-3, -8, and -9 triple knockout (TKO) mice. (**A**) Immunoblots of bone marrow-derived macrophage (BMM) lysates from WT and TKO mice (strain 2) probed for indicated proteins. (**B**) Quantification of Lysotracker puncta per cell after 30 min 0.5 mM LLOMe treatment followed by 30 min washout in WT or TKO BMMs. (**C**) Same as in (B) but 3 h 0.5 mM LLOMe treatment. Figures represent two independent experiments (B, C). Error bars represent SD from four technical replicates, and *p<0.05., ****p<0.0001 by unpaired t-test.

**Fig S3.**
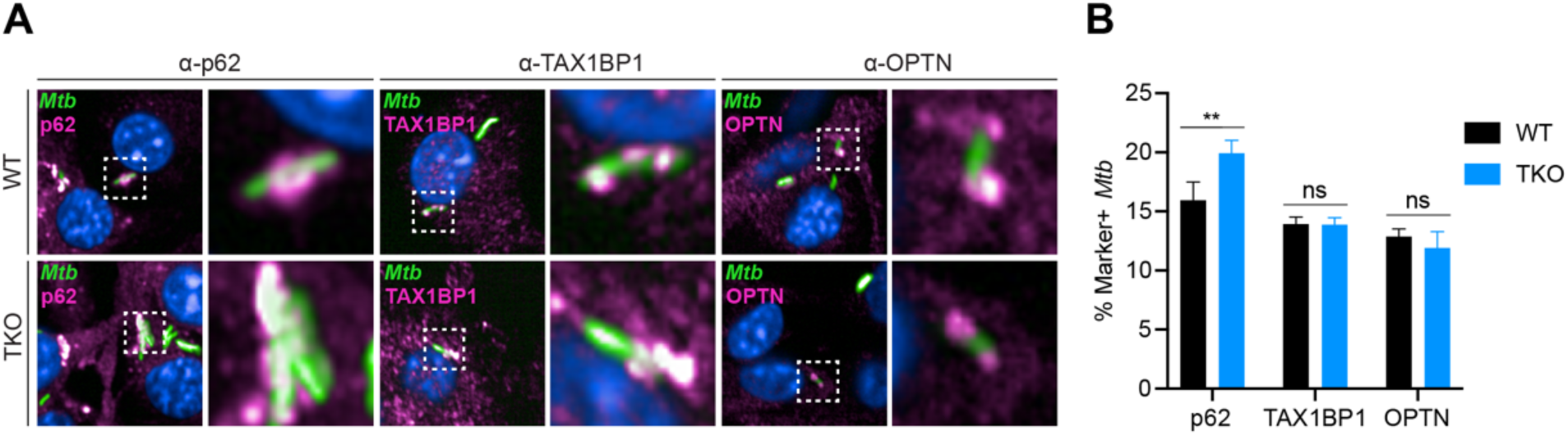
Autophagy receptor recruitment to *Mtb*. (**A**) Confocal microscopy of WT or TKO BMMs infected with *Mtb-*GFP (MOI = 2) 8 hours post-infection and immunostained for p62, TAX1BP1, or OPTN. (**B**) Quantification of (A) for *Mtb-*GFP colocalization with indicated markers. Figures are representative of two independent experiments. Error bars represent SD from four technical replicates, and **p<0.01 by unpaired t-test.

**Fig S4.**
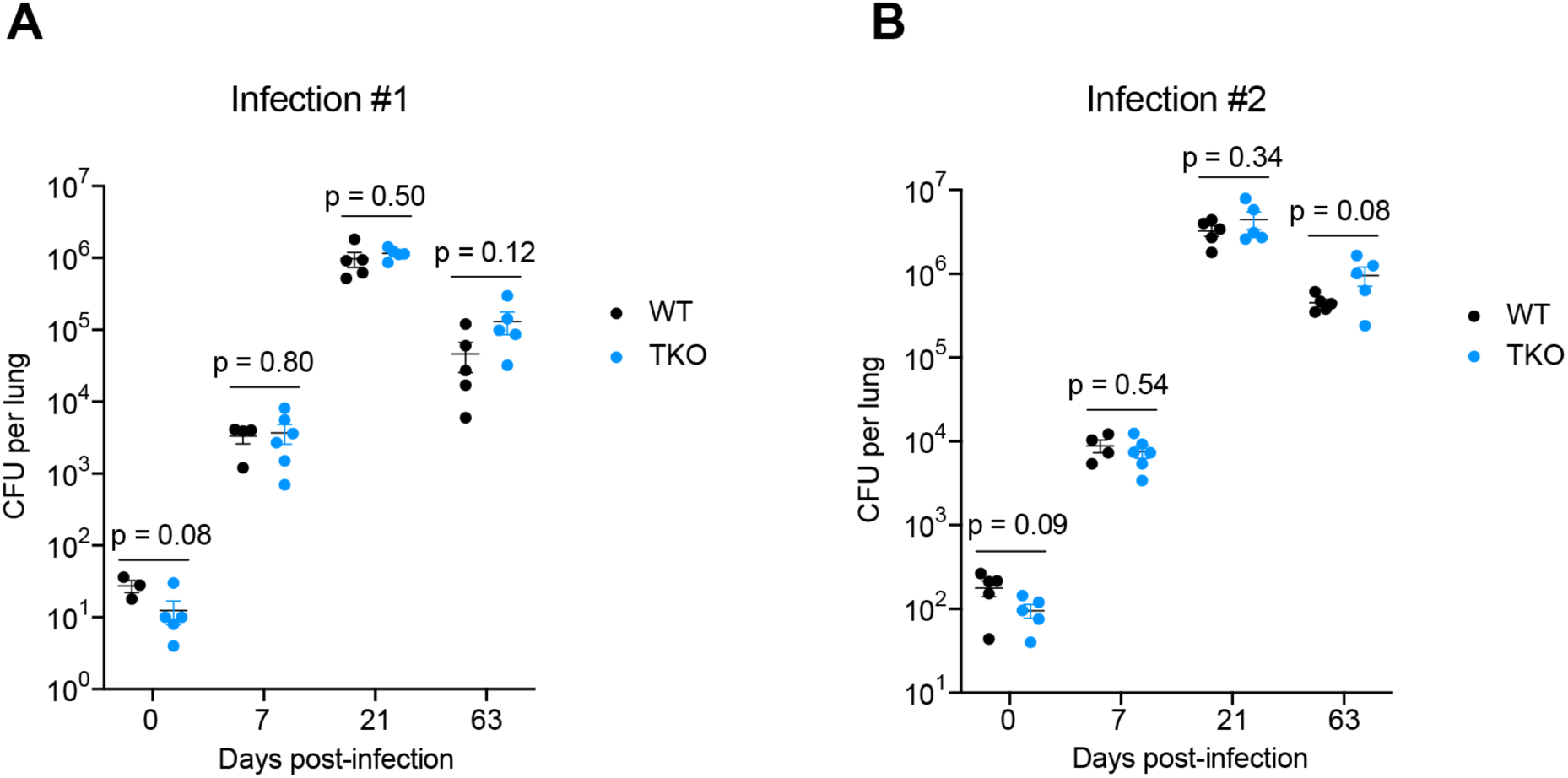
Lung CFU data for *Mtb* aerosol infections. WT and TKO female mice were aerosol infected with (**A**) ∼18 CFUs or (**B**) ∼136 CFUs of *Mtb* Erdman and bacterial loads in lungs were enumerated by plating for CFU at indicated time points. *n* = 3-5 mice per genotype. Figures represent one experiment. Bars represent the mean, error bars represent SEM, and p-values were determined by unpaired t-test.

**Fig S5.**
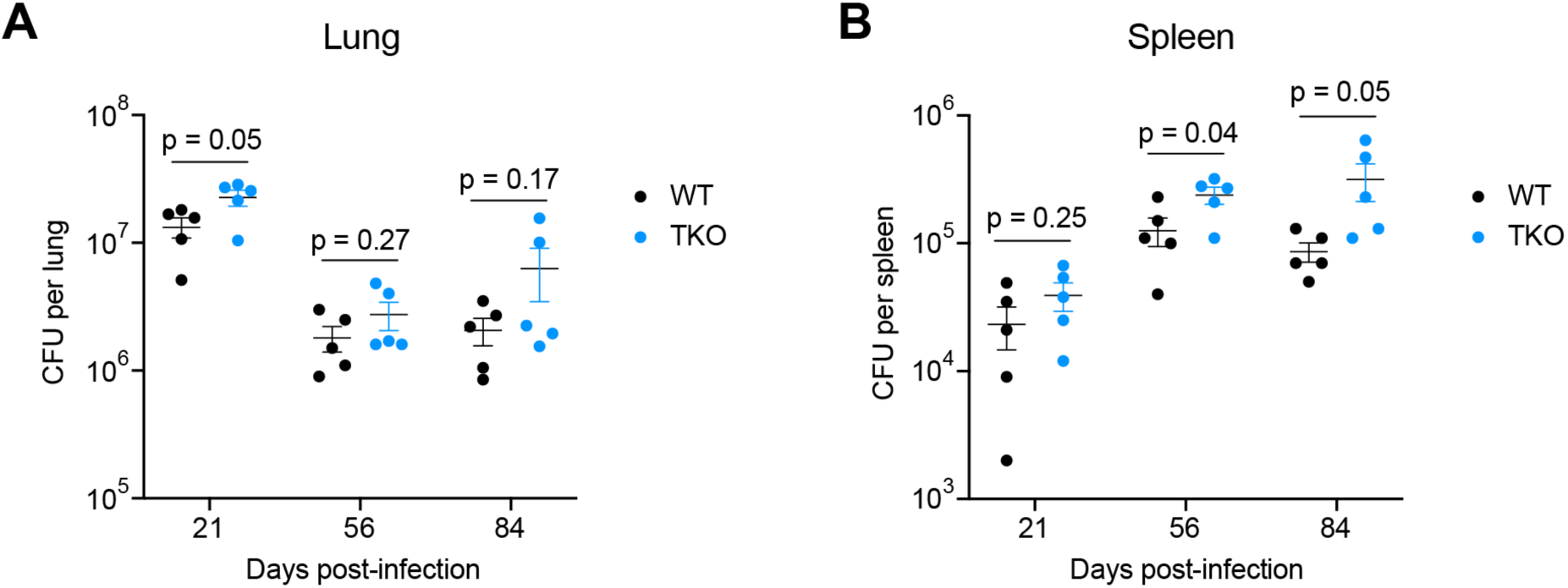
Membrane damage sensing galectins are partially required for control of *Mtb* in a high-dose aerosol infection. (**A**) WT and TKO male mice were aerosol infected with ∼450 CFUs of *Mtb* Erdman and bacterial loads in lungs were enumerated by plating for CFU at indicated time points. *n* = 5 mice per genotype. (**B**) Same as in (A) but bacterial loads in spleens. Figures represent one experiment. Bars represent the mean, error bars represent SEM, and p-values were determined by unpaired t-test.

**Fig S6.**
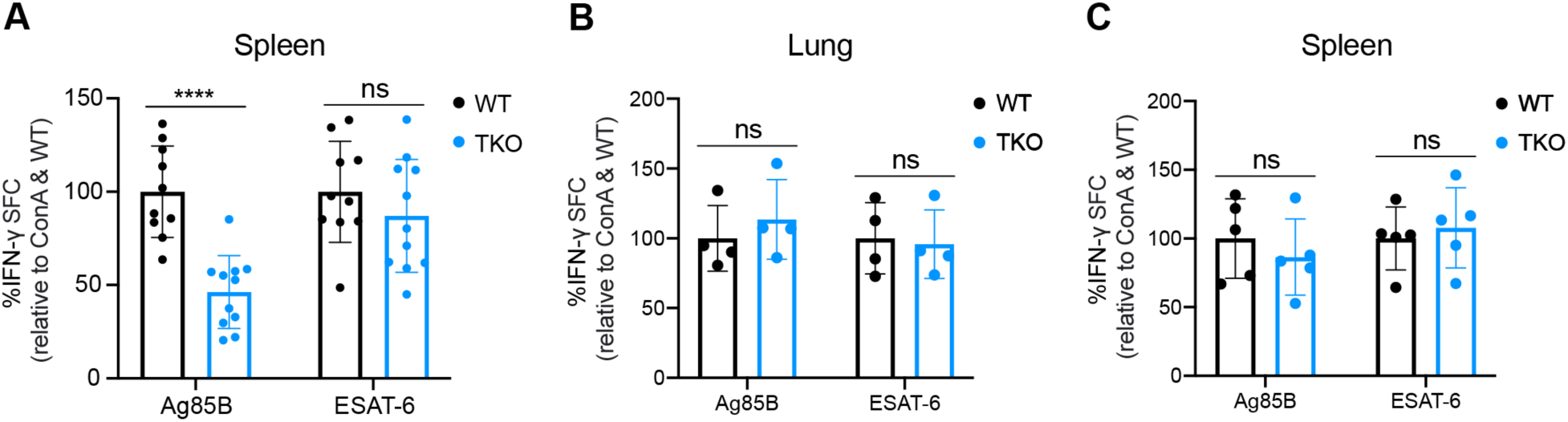
TKO mice exhibit defective splenic Ag85B-specific T cell responses. (**A**) WT and TKO mice were aerosol infected with ∼100 CFUs of *Mtb* Erdman and splenocytes were isolated for IFN-γ ELISpot to enumerate *Mtb*-specific T cells*. Mtb*-specific spot-forming cells (SFC) responding to re-stimulation in vitro with *Mtb* peptides are shown for each condition relative to Concanavalin A (ConA) total T cells and WT. *n* = 10-11 mice per genotype. (**B**) Same as in (A) but ELISpot performed with lung single cell suspension. *n* = 4 mice per genotype. (**C**) ELISpot on splenocytes following high-dose aerosol inoculation (∼800 CFUs). *n* = 5 mice per genotype. Figures represent data from three (A) independent experiments or one (B, C) independent experiment. Error bars represent SD, and ****p<0.0001 by unpaired t-test.

## References

1. Thakur A, Mikkelsen H, Jungersen G. Intracellular Pathogens: Host Immunity and Microbial Persistence Strategies. J Immunol Res. 2019;2019: 13565401. doi:10.1155/2019/1356540

2. Price J V, Vance RE. The macrophage paradox. Immunity. 2014;41: 685–693. doi:10.1016/j.immuni.2014.10.015

3. Watson RO, Bell SL, MacDuff DA, Kimmey JM, Diner EJ, Olivas J, et al. The Cytosolic Sensor cGAS Detects Mycobacterium tuberculosis DNA to Induce Type I Interferons and Activate Autophagy. Cell Host Microbe. 2015;17: 811–819. doi:10.1016/j.chom.2015.05.004

4. Kofoed EM, Vance RE. Innate immune recognition of bacterial ligands by NAIPs determines inflammasome specificity. Nature. 2011;477: 592–595. doi:10.1038/nature10394

5. Iwasaki A, Medzhitov R. Control of adaptive immunity by the innate immune system. Nat Immunol. 2015;16: 343–353. doi:10.1038/ni.3123

6. Ravesloot-Chávez MM, Van Dis E, Stanley SA. The Innate Immune Response to Mycobacterium tuberculosis Infection. Annu Rev Immunol. 2021;39: 611–637. doi:10.1146/annurev-immunol-093019-010426

7. Paz I, Sachse M, Dupont N, Mounier J, Cederfur C, Enninga J, et al. Galectin-3, a marker for vacuole lysis by invasive pathogens. Cell Microbiol. 2010;12: 530–544. doi:10.1111/j.1462-5822.2009.01415.x

8. Dupont N, Lacas-Gervais S, Bertout J, Paz I, Freche B, Van Nhieu GT, et al. Shigella phagocytic vacuolar membrane remnants participate in the cellular response to pathogen invasion and are regulated by autophagy. Cell Host Microbe. 2009;6: 137–149. doi:10.1016/j.chom.2009.07.005

9. Thurston TLM, Wandel MP, von Muhlinen N, Foeglein A, Randow F. Galectin 8 targets damaged vesicles for autophagy to defend cells against bacterial invasion. Nature. 2012;482: 414–418. doi:10.1038/nature10744

10. Rabinovich GA, Liu F-T, Hirashima M, Anderson A. An emerging role for galectins in tuning the immune response: lessons from experimental models of inflammatory disease, autoimmunity and cancer. Scand J Immunol. 2007;66: 143–158. doi:10.1111/j.1365-3083.2007.01986.x

11. Rabinovich GA, Toscano MA. Turning “sweet” on immunity: galectin-glycan interactions in immune tolerance and inflammation. Nat Rev Immunol. 2009;9: 338–352. doi:10.1038/nri2536

12. Cummings RD, Liu F-T. Galectins. In: Varki A, Cummings RD, Esko JD, Freeze HH, Stanley P, Bertozzi CR, et al., editors. Cold Spring Harbor (NY); 2009.

13. Jayaraman P, Sada-Ovalle I, Beladi S, Anderson AC, Dardalhon V, Hotta C, et al. Tim3 binding to galectin-9 stimulates antimicrobial immunity. J Exp Med. 2010;207: 2343– 2354. doi:10.1084/jem.20100687

14. Maejima I, Takahashi A, Omori H, Kimura T, Takabatake Y, Saitoh T, et al. Autophagy sequesters damaged lysosomes to control lysosomal biogenesis and kidney injury. EMBO J. 2013;32: 2336–2347. doi:10.1038/emboj.2013.171

15. Chauhan S, Kumar S, Jain A, Ponpuak M, Mudd MH, Kimura T, et al. TRIMs and Galectins Globally Cooperate and TRIM16 and Galectin-3 Co-direct Autophagy in Endomembrane Damage Homeostasis. Dev Cell. 2016;39: 13–27. doi:10.1016/j.devcel.2016.08.003

16. Jia J, Abudu YP, Claude-Taupin A, Gu Y, Kumar S, Choi SW, et al. Galectins Control mTOR in Response to Endomembrane Damage. Mol Cell. 2018;70: 120–135.e8. doi:10.1016/j.molcel.2018.03.009

17. Jia J, Claude-Taupin A, Gu Y, Choi SW, Peters R, Bissa B, et al. Galectin-3 Coordinates a Cellular System for Lysosomal Repair and Removal. Dev Cell. 2020;52: 69–87.e8. doi:10.1016/j.devcel.2019.10.025

18. Jia J, Bissa B, Brecht L, Allers L, Choi SW, Gu Y, et al. AMPK, a Regulator of Metabolism and Autophagy, Is Activated by Lysosomal Damage via a Novel Galectin-Directed Ubiquitin Signal Transduction System. Mol Cell. 2020;77: 951–969.e9. doi:10.1016/j.molcel.2019.12.028

19. Wong K-W, Jacobs WRJ. Critical role for NLRP3 in necrotic death triggered by Mycobacterium tuberculosis. Cell Microbiol. 2011;13: 1371–1384. doi:10.1111/j.1462-5822.2011.01625.x

20. Bell SL, Lopez KL, Cox JS, Patrick KL, Watson RO. Galectin-8 Senses Phagosomal Damage and Recruits Selective Autophagy Adapter TAX1BP1 To Control Mycobacterium tuberculosis Infection in Macrophages. MBio. 2021;12: e0187120. doi:10.1128/mBio.01871-20

21. Feeley EM, Pilla-Moffett DM, Zwack EE, Piro AS, Finethy R, Kolb JP, et al. Galectin-3 directs antimicrobial guanylate binding proteins to vacuoles furnished with bacterial secretion systems. Proc Natl Acad Sci U S A. 2017;114: E1698–E1706. doi:10.1073/pnas.1615771114

22. Pelletier I, Hashidate T, Urashima T, Nishi N, Nakamura T, Futai M, et al. Specific recognition of Leishmania major poly-beta-galactosyl epitopes by galectin-9: possible implication of galectin-9 in interaction between L. major and host cells. J Biol Chem. 2003;278: 22223–22230. doi:10.1074/jbc.M302693200

23. Stowell SR, Arthur CM, Dias-Baruffi M, Rodrigues LC, Gourdine J-P, Heimburg-Molinaro J, et al. Innate immune lectins kill bacteria expressing blood group antigen. Nat Med. 2010;16: 295–301. doi:10.1038/nm.2103

24. Quattroni P, Li Y, Lucchesi D, Lucas S, Hood DW, Herrmann M, et al. Galectin-3 binds Neisseria meningitidis and increases interaction with phagocytic cells. Cell Microbiol. 2012;14: 1657–1675. doi:10.1111/j.1462-5822.2012.01838.x

25. Wu X, Wu Y, Zheng R, Tang F, Qin L, Lai D, et al. Sensing of mycobacterial arabinogalactan by galectin-9 exacerbates mycobacterial infection. EMBO Rep. 2021;22: e51678. doi:10.15252/embr.202051678

26. Budzik JM, Swaney DL, Jimenez-Morales D, Johnson JR, Garelis NE, Repasy T, et al. Dynamic post-translational modification profiling of Mycobacterium tuberculosis-infected primary macrophages. Elife. 2020;9. doi:10.7554/eLife.51461

27. Johnson JR, Parry T, Repasy T, Geiger KM, Verschueren E, Budzik JM, et al. Comparative analysis of macrophage post-translational modifications during intracellular bacterial pathogen infection. bioRxiv. 2020; 2020.05.27.116772. doi:10.1101/2020.05.27.116772

28. Mizushima N. Autophagy: process and function. Genes Dev. 2007;21: 2861–2873. doi:10.1101/gad.1599207

29. Levine B, Mizushima N, Virgin HW. Autophagy in immunity and inflammation. Nature. 2011;469: 323–335. doi:10.1038/nature09782

30. Manzanillo PS, Ayres JS, Watson RO, Collins AC, Souza G, Rae CS, et al. The ubiquitin ligase parkin mediates resistance to intracellular pathogens. Nature. 2013;501: 512–516. doi:10.1038/nature12566

31. Franco LH, Nair VR, Scharn CR, Xavier RJ, Torrealba JR, Shiloh MU, et al. The Ubiquitin Ligase Smurf1 Functions in Selective Autophagy of Mycobacterium tuberculosis and Anti-tuberculous Host Defense. Cell Host Microbe. 2017;21: 59–72. doi:10.1016/j.chom.2016.11.002

32. Stolz A, Ernst A, Dikic I. Cargo recognition and trafficking in selective autophagy. Nat Cell Biol. 2014;16: 495–501. doi:10.1038/ncb2979

33. Watson RO, Manzanillo PS, Cox JS. Extracellular M. tuberculosis DNA targets bacteria for autophagy by activating the host DNA-sensing pathway. Cell. 2012;150: 803–815. doi:10.1016/j.cell.2012.06.040

34. Braverman J, Sogi KM, Benjamin D, Nomura DK, Stanley SA. HIF-1α Is an Essential Mediator of IFN-γ-Dependent Immunity to Mycobacterium tuberculosis. J Immunol. 2016;197: 1287–1297. doi:10.4049/jimmunol.1600266

35. Ren T, Zamboni DS, Roy CR, Dietrich WF, Vance RE. Flagellin-deficient Legionella mutants evade caspase-1- and Naip5-mediated macrophage immunity. PLoS Pathog. 2006;2: e18. doi:10.1371/journal.ppat.0020018

36. Mitchell G, Cheng MI, Chen C, Nguyen BN, Whiteley AT, Kianian S, et al. Listeria monocytogenes triggers noncanonical autophagy upon phagocytosis, but avoids subsequent growth-restricting xenophagy. Proc Natl Acad Sci U S A. 2018;115: E210– E217. doi:10.1073/pnas.1716055115

37. Cooper AM. Cell-mediated immune responses in tuberculosis. Annu Rev Immunol.2009;27: 393–422. doi:10.1146/annurev.immunol.021908.132703

38. Sakai S, Kauffman KD, Schenkel JM, McBerry CC, Mayer-Barber KD, Masopust D, et al. Cutting edge: control of Mycobacterium tuberculosis infection by a subset of lung parenchyma-homing CD4 T cells. J Immunol. 2014;192: 2965–2969. doi:10.4049/jimmunol.1400019

39. Tattoli I, Sorbara MT, Yang C, Tooze SA, Philpott DJ, Girardin SE. Listeria phospholipases subvert host autophagic defenses by stalling pre-autophagosomal structures. EMBO J. 2013;32: 3066–3078. doi:10.1038/emboj.2013.234

40. Mitchell G, Ge L, Huang Q, Chen C, Kianian S, Roberts MF, et al. Avoidance of autophagy mediated by PlcA or ActA is required for Listeria monocytogenes growth in macrophages. Infect Immun. 2015;83: 2175–2184. doi:10.1128/IAI.00110-15

41. Choy A, Dancourt J, Mugo B, O’Connor TJ, Isberg RR, Melia TJ, et al. The Legionella effector RavZ inhibits host autophagy through irreversible Atg8 deconjugation. Science. 2012;338: 1072–1076. doi:10.1126/science.1227026

42. Conway KL, Kuballa P, Song J-H, Patel KK, Castoreno AB, Yilmaz OH, et al. Atg16l1 is required for autophagy in intestinal epithelial cells and protection of mice from Salmonella infection. Gastroenterology. 2013;145: 1347–1357. doi:10.1053/j.gastro.2013.08.035

43. Kobayashi KS, Chamaillard M, Ogura Y, Henegariu O, Inohara N, Nuñez G, et al. Nod2-dependent regulation of innate and adaptive immunity in the intestinal tract. Science. 2005;307: 731–734. doi:10.1126/science.1104911

44. Castillo EF, Dekonenko A, Arko-Mensah J, Mandell MA, Dupont N, Jiang S, et al. Autophagy protects against active tuberculosis by suppressing bacterial burden and inflammation. Proc Natl Acad Sci U S A. 2012;109: E3168–76. doi:10.1073/pnas.1210500109

45. Kimmey JM, Huynh JP, Weiss LA, Park S, Kambal A, Debnath J, et al. Unique role for ATG5 in neutrophil-mediated immunopathology during M. tuberculosis infection. Nature. 2015;528: 565–569. doi:10.1038/nature16451

46. Dengjel J, Schoor O, Fischer R, Reich M, Kraus M, Müller M, et al. Autophagy promotes MHC class II presentation of peptides from intracellular source proteins. Proc Natl Acad Sci U S A. 2005;102: 7922–7927. doi:10.1073/pnas.0501190102

47. Levine B, Deretic V. Unveiling the roles of autophagy in innate and adaptive immunity. Nat Rev Immunol. 2007;7: 767–777. doi:10.1038/nri2161

48. Paludan C, Schmid D, Landthaler M, Vockerodt M, Kube D, Tuschl T, et al. Endogenous MHC class II processing of a viral nuclear antigen after autophagy. Science. 2005;307: 593–596. doi:10.1126/science.1104904

49. Lee HK, Mattei LM, Steinberg BE, Alberts P, Lee YH, Chervonsky A, et al. In vivo requirement for Atg5 in antigen presentation by dendritic cells. Immunity. 2010;32: 227– 239. doi:10.1016/j.immuni.2009.12.006

50. Saini NK, Baena A, Ng TW, Venkataswamy MM, Kennedy SC, Kunnath-Velayudhan S, et al. Suppression of autophagy and antigen presentation by Mycobacterium tuberculosis PE_PGRS47. Nat Microbiol. 2016;1: 16133. doi:10.1038/nmicrobiol.2016.133

51. Moguche AO, Shafiani S, Clemons C, Larson RP, Dinh C, Higdon LE, et al. ICOS and Bcl6-dependent pathways maintain a CD4 T cell population with memory-like properties during tuberculosis. J Exp Med. 2015;212: 715–728. doi:10.1084/jem.20141518

52. Rogerson BJ, Jung Y-J, LaCourse R, Ryan L, Enright N, North RJ. Expression levels of Mycobacterium tuberculosis antigen-encoding genes versus production levels of antigen-specific T cells during stationary level lung infection in mice. Immunology. 2006;118: 195–201. doi:10.1111/j.1365-2567.2006.02355.x

53. Moguche AO, Musvosvi M, Penn-Nicholson A, Plumlee CR, Mearns H, Geldenhuys H, et al. Antigen Availability Shapes T Cell Differentiation and Function during Tuberculosis. Cell Host Microbe. 2017;21: 695–706.e5. doi:10.1016/j.chom.2017.05.012

54. Sani M, Houben ENG, Geurtsen J, Pierson J, de Punder K, van Zon M, et al. Direct visualization by cryo-EM of the mycobacterial capsular layer: a labile structure containing ESX-1-secreted proteins. PLoS Pathog. 2010;6: e1000794. doi:10.1371/journal.ppat.1000794

55. Daffé M. The cell envelope of tubercle bacilli. Tuberculosis (Edinb). 2015;95 Suppl 1: S155–8. doi:10.1016/j.tube.2015.02.024

56. Beatty WL, Rhoades ER, Ullrich HJ, Chatterjee D, Heuser JE, Russell DG. Trafficking and release of mycobacterial lipids from infected macrophages. Traffic. 2000;1: 235–247. doi:10.1034/j.1600-0854.2000.010306.x

57. Penn BH, Netter Z, Johnson JR, Von Dollen J, Jang GM, Johnson T, et al. An Mtb-Human Protein-Protein Interaction Map Identifies a Switch between Host Antiviral and Antibacterial Responses. Mol Cell. 2018;71: 637–648.e5. doi:10.1016/j.molcel.2018.07.010

58. Sogi KM, Lien KA, Johnson JR, Krogan NJ, Stanley SA. The Tyrosine Kinase Inhibitor Gefitinib Restricts Mycobacterium tuberculosis Growth through Increased Lysosomal Biogenesis and Modulation of Cytokine Signaling. ACS Infect Dis. 2017;3: 564–574. doi:10.1021/acsinfecdis.7b00046

59. Gonçalves A V, Margolis SR, Quirino GFS, Mascarenhas DPA, Rauch I, Nichols RD, et al. Gasdermin-D and Caspase-7 are the key Caspase-1/8 substrates downstream of the NAIP5/NLRC4 inflammasome required for restriction of Legionella pneumophila. PLoS Pathog. 2019;15: e1007886. doi:10.1371/journal.ppat.1007886

60. Reniere ML, Whiteley AT, Portnoy DA. An In Vivo Selection Identifies Listeria monocytogenes Genes Required to Sense the Intracellular Environment and Activate Virulence Factor Expression. PLoS Pathog. 2016;12: e1005741. doi:10.1371/journal.ppat.1005741

61. Coers J, Vance RE, Fontana MF, Dietrich WF. Restriction of Legionella pneumophila growth in macrophages requires the concerted action of cytokine and Naip5/Ipaf signalling pathways. Cell Microbiol. 2007;9: 2344–2357. doi:10.1111/j.1462-5822.2007.00963.x

62. Wang C, Mei L. In utero electroporation in mice. Methods Mol Biol. 2013;1018: 151– 163. doi:10.1007/978-1-62703-444-9_15

63. Van Dis E, Sogi KM, Rae CS, Sivick KE, Surh NH, Leong ML, et al. STING-Activating Adjuvants Elicit a Th17 Immune Response and Protect against Mycobacterium tuberculosis Infection. Cell Rep. 2018;23: 1435–1447. doi:10.1016/j.celrep.2018.04.003

